# A Binuclear [Cu_2_S_2_] Cluster Supports Functional Asymmetry in Dark-Operative Protochlorophyllide Reductase

**DOI:** 10.64898/2026.11.03.763803

**Authors:** Kewei Zhao, Rajnandani Kashyap, Macon J. Abernathy, Edwin Antony, Ritimukta Sarangi

## Abstract

Many photosynthetic bacteria rely on the nitrogenase-like dark-operative protochlorophyllide oxidoreductase (DPOR) to sustain (bacterio)chlorophyll synthesis in the dark. DPOR couples ATP hydrolysis to electron transfer from a reducing component (L-protein) to a catalytic component (NB-protein) that reduces protochlorophyllide. NB-protein is an α_2_β_2_ heterotetramer, and each heterodimeric half contains an active site and a [Fe_4_S_4_] cluster. Although the two halves are identical, they react sequentially rather than in parallel. How the two halves communicate has remained unknown. Here we show, using Cu K-edge x-ray absorption spectroscopy and cryo-EM, that the NB-protein tetramer interface holds a binuclear, sulfur-bridged [Cu_2_S_2_] cluster. Each Cu is reduced and three-coordinate, with short Cu-S and Cu-Cu distances (2.26 and 2.62 Å). Mutagenesis excludes the nearest cysteine as the bridging ligand, supporting inorganic sulfide as the most likely sulfur donor. To our knowledge, this is the first sulfide-bridged dicopper cluster identified in a protein. The cluster is conserved in three purple non-sulfur bacteria and connects the BchB C-terminal helices that contact opposing active sites. Loss of Cu abolishes DPOR activity and asymmetric L-protein binding without disturbing overall fold. We propose that the cluster relays substrate-binding information between active sites, suggesting a metal-mediated mechanism for cooperativity in nitrogenase-like enzymes.

## Introduction

Biochemical processes that facilitate carbon fixation in photosynthetic organisms are essential to the global carbon cycle and represent potential blueprints for bio-energy technologies^1^. Dark-operative protochlorophyllide Oxidoreductase (DPOR) is a key enzyme in the bacteriochlorophyll biosynthetic pathway that catalyzes the stereospecific reduction of protochlorophyllide (Pchlide) to chlorophyllide, enabling photosynthetic organisms to synthesize chlorophyll in the absence of light^2–4^. It belongs to the nitrogenase-like superfamily and resembles nitrogenase in architecture and mechanism, both harnessing ATP binding and hydrolysis to drive electron transfer through a series of metal clusters^5,6^. DPOR consists of two components: the homodimeric reducing component BchL protein (L-protein) contains an [Fe_4_S_4_]^2+^ cluster and an ATP-binding site, and the heterotetrameric catalytic component BchN-BchB protein (NB-protein) contains one [Fe_4_S_4_]^2+^ cluster and one Pchlide-binding site in each half, at positions structurally analogous to the P-cluster and the FeMo-cofactor of nitrogenase.

Although the DPOR complex has a two-fold symmetry, it does not operate symmetrically^7–9^. The current mechanistic model proposes that Pchlide binding triggers electron transfer that proceeds sequentially rather than in parallel across the two halves, and disrupting electron transfer in one half abolishes turnover of the entire complex^7^. Therefore, some structural elements must relay the substrate-binding and turnover states of one active site to the other. Our recent cryogenic electron microscopy (cryo-EM) analysis of *Rhodobacter sphaeroides* DPOR identified two Cu ions at the center of the NB-protein as a candidate for this communication role^8^. The Cu site bridges the C-terminal α-helices of the two BchB subunits, which cross at the tetramer interface and each contacts the Pchlide site of the opposing half. Alanine substitution of His-404 and Met-408 adjacent to the Cu site eliminated both Cu incorporation and Pchlide reduction activity, indicating that Cu is required for catalysis. However, the chemistry of this site, e.g. the oxidation state and coordination structure of Cu, remained undefined in the cryo-EM structure.

Here we address these questions using Cu K-edge x-ray absorption spectroscopy (XAS), cryo-EM, and site-directed mutagenesis. We show that the Cu site is not a conventional His/Met Cu center but a compact binuclear [Cu_2_S_2_] cluster with three-coordinate Cu(I), and it is conserved in NB-proteins from three purple non-sulfur bacteria. The bridging sulfur ligands are most consistent with inorganic sulfides. The structural resemblance to Cu-trafficking clusters rather than to catalytic Cu centers, together with its placement at the tetramer interface, implies that the cluster mediates allosteric communication by coupling the two halves of DPOR. In support of this model, a cryo-EM structure of the Cu-binding mutant showcases a loss in asymmetry and allostery between the two halves.

## Results

### A central Cu site bridges the two halves of NB-protein

In our previous cryo-EM study of *Rhodobacter sphaeroides* DPOR, two Cu ions were positioned at the center of NB-protein, where two helices from opposing BchB subunits cross at the tetramer interface^8^. The Cu ions were modeled near His-404 and Met-408 residues, and alanine substitution of these residues abolished Cu incorporation and Pchlide reduction activity. To examine whether loss of Cu affects the structure of NB-protein, we analyzed the H404A-M408A double mutant with cryo-EM in the presence of L-protein and ATP. The mutant retains the heterotetrameric architecture and forms a complex with two L-proteins (Fig. 1a). These data show that the loss in activity does not arise from a loss in secondary structure or a failure to engage the L-protein. However, unlike the wild-type protein (Fig. 1b), two L-proteins are bound to the NB-protein tetramer in the presence of ATP and Pchlide (turnover conditions). This symmetric engagement of L-proteins on both sides resembles the trapped-state of the DPOR complex that is observed in the presence of ADP-AlF_3_ (Fig. 1c). The central metal density observed in the wild-type protein is also absent, and the mutant contains a hollow pocket at the same position (Fig. 1a). These data support that the Cu site is located at the interface of two halves of the NB-protein, and His-404 and Met-408 are required for its incorporation and relevant to the allostery within DPOR.

**Figure 1.**
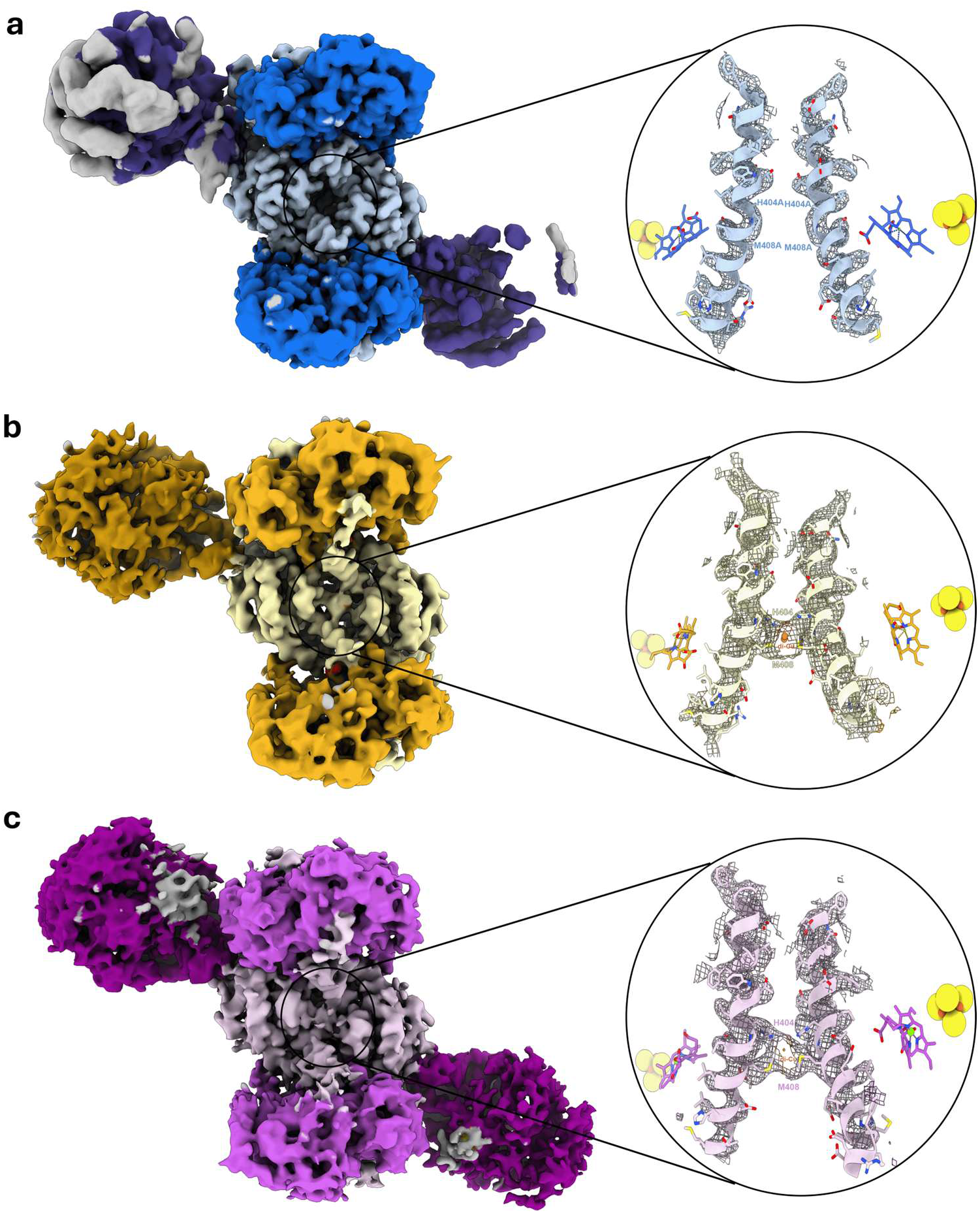
Cryo-EM structure of the DPOR H404A-M408A mutant complex during turnover. **a,** Cryo-EM reconstruction of *Rhodobacter sphaeroides* DPOR containing the BchB H404A-M408A double mutant under turnover conditions, in the presence of ATP and Pchlide. The mutant retains the heterotetrameric NB-protein architecture and binds two L-protein dimers on opposite halves of the complex, but lacks the central density assigned to Cu in the wild-type protein. **b,** Wild-type DPOR under turnover conditions, showing asymmetric binding of a single L-protein dimer and retention of the central Cu site. **c,** Wild-type DPOR trapped with the transition-state analog ADP-AlF_3_, showing two bound L-protein dimers, as observed for the double mutant under turnover conditions. Insets show the opposing BchB C-terminal α-helices at the NB-protein interface, with atomic models superimposed on the cryo-EM density (mesh), highlighting the absence of central Cu density in the double mutant.

A long helix in BchB (residues 398–420) interacts in *trans* with the active site and the bound Pchlide molecule at the opposing half of the NB-tetramer. In the ADP-AlF_3_-bound transition state structure of DPOR, this helix transitions between bent versus extended conformations, suggesting a role in substrate authentication ^8^. In the DPOR double mutant, while the asymmetry is lost and two L-proteins are bound to the NB-protein, we observe no structural transitions in this helix. These findings suggest that the Cu cluster might be responsible for relaying information about the substrate occupancy and redox state.

The coordination environment of the Cu ions was not fully characterized by cryo-EM: the modeled Cu-ligand distances (> 3 Å) are longer than expected for typical Cu-N and Cu-S bonds, thus it is unclear whether His-404 and Met-408 are coordinated to the Cu ions. The electron paramagnetic resonance (EPR) silence of native NB-protein (purified in the presence of excess reductant) precludes electronic structure characterization using conventional methods^8^. Here, we employed Cu K-edge high-energy-resolution fluorescence-detected (HERFD) and conventional XAS to evaluate the geometric and electronic structure of the Cu site.

### X-ray spectroscopy reveals a binuclear Cu(I)-sulfur cluster

Consistent with the absence of central metal density in the cryo-EM structure, the H404A-M408A mutant exhibited little Cu Kα fluorescence signal, with less than 3% of the count rate observed for the wild-type NB-protein. EPR spin-quantification of H_2_O_2_-oxidized samples shows a Cu-to-protein ratio of 1.7:1 for the wild-type sample and only background-level Cu in the double mutant (Fig. S1a). ICP-MS measurements similarly show 1.8 Cu per wild-type protein (Table S1) and little Cu in the mutant. This confirms that His-404 and Met-408 are required for native Cu incorporation.

Cu K-edge XAS of the wild-type (WT) NB-protein of *R. sphaeroides* were measured to define the chemical structure of the central Cu site. The normalized spectrum shows an absorption edge at ∼8981.5 eV, consistent with a Cu center in the +1 oxidation state (Fig. 2a). No pre-edge feature is observed, as expected for closed-shell 3d^10^ Cu(I) (consistent with its EPR silence)^8^. The spectrum exhibits a pronounced rising-edge feature near ∼8983 eV. This feature has been assigned to Cu 1s→4p transitions, and its intensity and energy position have been shown to be diagnostic of the local coordination geometry around the Cu center^10^. The comparison with reference compounds shows that NB-protein is consistent with a three-coordinate Cu(I) site.^10^

**Figure 2.**
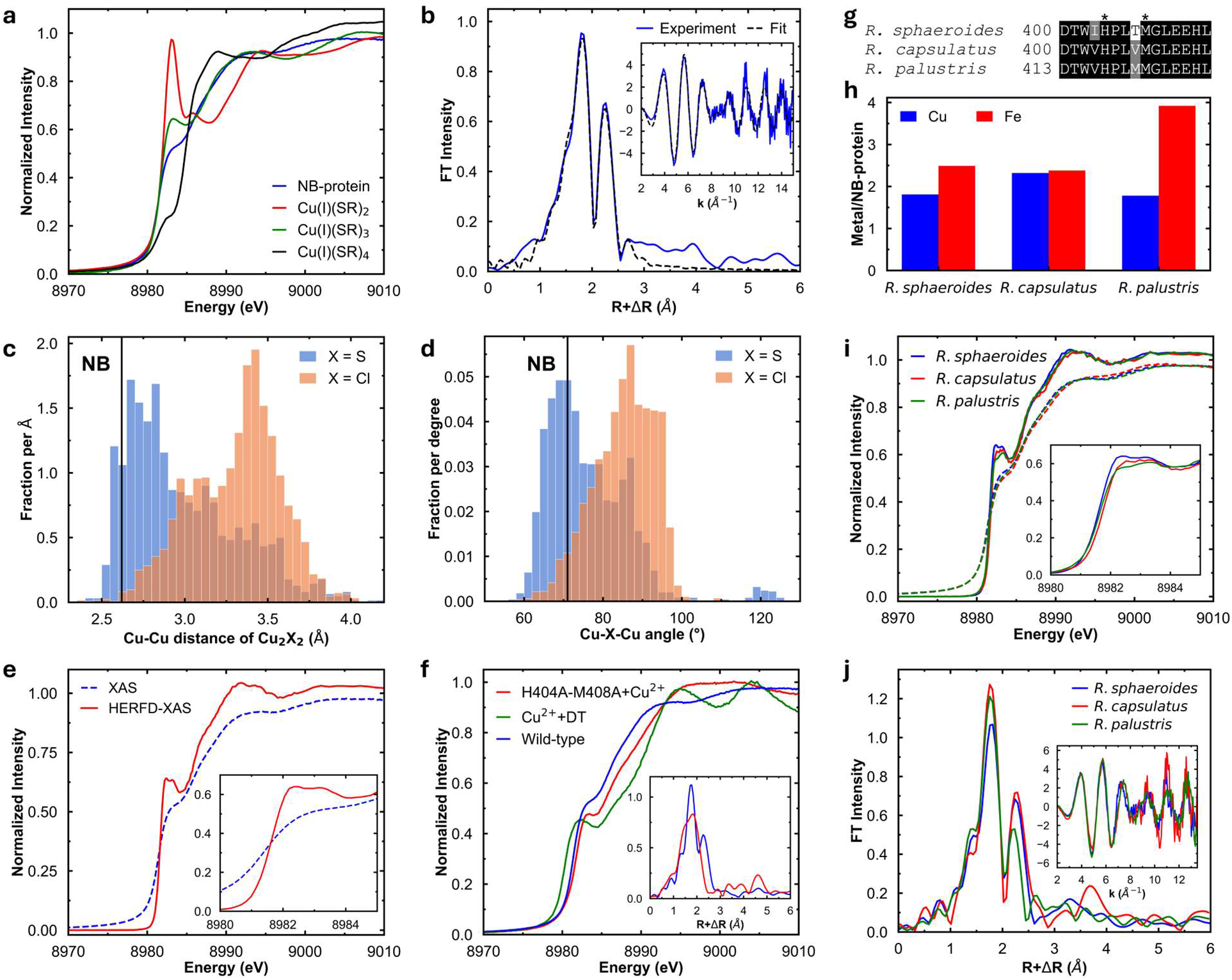
Characterization of the Cu site in NB-protein. **a,** Cu K-edge XAS of NB-protein and reference Cu(I)-S compounds with varying coordination numbers. **b**, Cu K-edge EXAFS of NB-protein (inset) and the Fourier transform (FT). The fit is shown in dashed lines. **c**, Distribution of Cu-Cu distances in Cu_2_X_2_ cores surveyed from the Cambridge Structural Database (n = 1244 and 2489 for X = S and Cl, respectively). The black vertical line marks the EXAFS-derived Cu-Cu distance of the NB cluster. **d,** Distribution of Cu-X-Cu angles in the same Cu_2_X_2_ cores. The Cu-S-Cu angle of the NB cluster (black line) was calculated from the EXAFS-derived Cu-Cu and Cu-S distances, assuming a symmetric Cu_2_S_2_ rhombic core. **e,** Cu K-edge HERFD-XAS of NB-protein. The inset zooms in on the rising edge. **f,** Cu K-edge XAS of the Cu-supplemented H404A-M408A double mutant and the Cu^2+^-dithionite control sample. The inset compares the FT of the EXAFS of the wild-type protein and Cu-supplemented mutant. **g,** sequence alignment of BchB proteins of three purple non-sulfur bacteria. The asterisks denote His-404 and Met-408. **h,** ICP-MS quantification of the metal-to-NB-protein ratio of three purple non-sulfur bacteria. **i,** Cu K-edge XAS (dashed lines) and HERFD-XAS (solid lines) of NB-protein from three purple non-sulfur bacteria. The inset zooms in on the rising edge. **j,** Cu K-edge EXAFS (inset) and FT of the three NB-proteins.

Cu K-edge extended x-ray absorption fine structure (EXAFS) of NB-protein were measured to k = 15 Å^-1^ (Fig. 2b). The spectrum exhibits two strong sinusoidal envelopes, with a distinctive beat between k = 8-9 Å^-1^. The Fourier transform (FT) is dominated by two features at R’ = 1.8 and 2.3 Å, respectively, indicating a well-ordered coordination environment with both first-shell ligation and a heavy, more distant scatterer. Attempts were made to fit the EXAFS data with a structural model derived from the cryo-EM structure, in which light-atom ligands, such as water and histidine, and the heavier methionine sulfur were included in the first and second shells, respectively. A Cu-Cu interaction was also modeled, starting at the cryo-EM distance. Since the wild-type cryo-EM structure was obtained at ∼2.70 Å resolution, several structural models using a binuclear Cu motif were tested. All models underestimate the low-k EXAFS amplitude and the first-shell FT intensity (best-fit model using the cryo-EM structure is shown in Fig. S2 and Table S2). The best-fits to the data instead are consistent with 0.5 Cu-N/O path at ∼1.95 Å, two Cu-S paths at ∼2.26 Å, one Cu-Cu path at ∼2.62 Å, and a longer, 0.5 or 1 Cu-S path at ∼2.89 Å (Fig. 2b and S3, Table S2).

The Cu-Cu path makes a major contribution to the FT feature at R’ = 2.3 Å, supporting a compact, doubly bridged binuclear Cu(I) site^11,12^. The 2.26 Å Cu-S distance is characteristic of thiolate or sulfide ligands^13–15^ and argues against strong Met-408 coordination. Met-408 may lead to the 2.89 Å component, however two Cu-O paths can also account for this feature with somewhat lower fit quality (Table S2). The ∼0.5 Cu-N coordination makes a small but real impact on the EXAFS fit (2% F-factor improvement), suggesting an asymmetric site with a single light atom ligand to one of the Cu centers^16–18^. This could be the histidine ligand or a water molecule. We note that a chloride-bridged dimer with a Cu-Cl at 2.26 Å would be indistinguishable from a Cu-S interaction. While NaCl was present in the samples, it is unlikely to form a reduced Cu(I)-Cu(I) dimer. A survey of the Cambridge Structural Database^19^ shows that Cu_2_Cl_2_ species have a mean Cu-Cu distance of ∼3.5 Å (Fig. 2c) and a greater bridging angle than the Cu_2_S_2_ species (Fig. 2d), therefore the NB cluster geometry does not support chlorine as the bridging ligand. The EXAFS model presented above is the minimal one that reproduces the data, and additional longer-distance shells can be accommodated but are not required.

UV absorption data of the NB-protein were measured and compared to that of the double mutant (Fig. S4). An increase in intensity at ∼260 nm is consistent with Cu-S charge transfer band in Cu(I)-S compounds^20^. Although the intensity difference is small, neither the [Fe_4_S_4_]^2+^ cluster nor dithionite, which absorb at 290 nm^21^ and 315 nm^22^, respectively, contribute appreciably in the 260 nm region. Together, our data (EXAFS, EPR, ICP-MS, cryo-EM and UV-vis) support the presence of a binuclear Cu-S cluster in which each Cu is three-coordinate, connected to the C-terminal α-helices through weak interactions.

The unequal Cu-ligand distances suggest a deviation from ideal trigonal coordination. To probe the local symmetry, we used Cu K-edge HERFD XAS to better resolve the symmetry-sensitive 1s→4p region. Here, the single feature observed in conventional XAS resolves into two peaks separated by ∼1 eV (Fig. 2e). Symmetry considerations indicate that out-of-plane displacement of Cu can induce 4s-4p mixing and explain this splitting^23^. In a recent report, time-dependent density functional theory (TD-DFT) was successfully used to quantitatively study HERFD-XAS of Cu(I) compounds^24^. A similar DFT/TD-DFT protocol was applied to a simplified trigonal [Cu(SCH_3_)_3_]^2-^ model. In the ideal trigonal planar geometry, the Cu 1s→4p transition is dominated by a single feature (Fig. S5). Displacing Cu out of the trigonal plane lowers the site symmetry, red-shifting the transition and splitting it into two. Molecular orbital analysis reveals that the two acceptor orbitals show mixed Cu 4s and 4p_z_ character with negligible 4p_x/y_ contribution (Table S3). Note that constraining the structure with asymmetric Cu-S distances also impacts the energies and relative intensities of these transitions (Fig. S5), thus the observed 1s→4p intensity represents an interplay of both factors and supports deviation from a trigonal planar geometry.

### His-404 and Met-408 are required for Cu cluster incorporation

Cu addition assays using the H404A-M408A double mutant were conducted to test whether the ordered Cu site of the WT protein could instead arise from adventitious Cu binding. Cu K-edge XAS data were collected on the double mutant treated with aqueous Cu^2+^ at a 1.8:1 Cu-to-protein ratio, which shows the Cu sites get fully reduced to Cu(I) in the presence of dithionite (Fig. 2f). The spectrum is distinct from that of the WT, indicating that the added Cu is coordinated differently to the mutant. EXAFS data reveal a site significantly disordered relative to the WT protein: strong low-k oscillations are present that attenuate rapidly beyond k = 8 Å^-1^, and the spectrum lacks the characteristic beat between k = 8-9 Å^-1^ (Fig. S1b). Best fits reveal a disordered Cu(I) environment with mixed N/O and S ligation (Table S2), likely reflecting a distribution of nonspecific protein-binding sites rather than a single, well-defined cluster. Therefore, Cu addition does not restore the wild-type active site structure in the double mutant.

### The Cu-S cluster is conserved across three bacterial NB-proteins

DPOR is found in diverse anaerobic photosynthetic bacteria, prompting the question: is the Cu-S cluster observed in *R. sphaeroides* NB-protein unique to this species, or is it conserved among DPORs? To address this, we heterologously expressed and purified NB-proteins of *Rhodobacter capsulatus* and *Rhodopseudomonas palustris* (Fig. S6a). The NB-proteins of the three purple non-sulfur bacteria share similar sequences, including conservation of the His-404 and Met-408 residues associated with Cu incorporation in *R. sphaeroides* (Fig. 2g). ICP-MS analysis detected approximately two Cu ions per NB-protein complex in both *R. capsulatus* and *R. palustris*, similar to the stoichiometry observed for *R. sphaeroides* NB-protein (Fig. 2h). These data indicate that Cu incorporation is not unique to *R. sphaeroides*.

Cu K-edge XAS reveals that the Cu sites in the three NB-proteins have similar oxidation states and coordination environments. The spectra for *R. capsulatus* and *R. palustris* align closely with those of *R. sphaeroides* (Fig. 2i). Conventional and HERFD-XAS spectra show similar rising-edge features assigned to the Cu(I) 1s→4p transition that are consistent with three-coordinate Cu(I), although modest differences in intensity suggest small variations in coordination geometry. EXAFS data for *R. capsulatus* and *R. palustris* exhibit the characteristic beat between k = 8-9 Å^-1^ and their Fourier transforms have the same two-shell pattern observed for *R. sphaeroides* NB-protein (Figs. 2j). Best fits for the EXAFS reveal very similar binuclear Cu-S cluster models (Fig. S7), and the derived Cu-S and Cu-Cu distances are consistent across species (Table S2) while the weaker Cu-N/O and Cu-S(Met) interactions show greater variation. Together, the sequence conservation, Cu stoichiometry, and XAS/EXAFS similarity support the presence of this binuclear cluster as a conserved feature of NB-proteins across multiple anaerobic bacteria. We also note that the anaerobic crystal structure of *R. capsulatus* NB-protein reports additional electron density at the tetramer interface (PDB 3AEQ), the position equivalent to the Cu site in our model, and the density was modeled as two water molecules^25^. Their refined B-factors (25.7 and 26.5 Å_2_) are significantly lower than surrounding protein atoms (mean ∼50 Å^2^), raising the possibility that the density instead arises from heavier atoms.

### Reducing conditions promote ordered Cu-S cluster assembly

The native NB-protein was prepared in the presence of 1.7 mM dithionite to maintain a reducing environment during expression and purification required for proper loading of the [Fe_4_S_4_]^2+^ clusters^26^. To test the impact of the reducing environment on the Cu cluster assembly, we prepared *R. sphaeroides* NB-protein in the absence of dithionite for ICP-MS and Cu K-edge XAS analysis. The Cu-to-protein ratio (Table S1) and Cu Kα fluorescence count rate (variation ∼15%) of the dithionite-free sample are comparable to those of the dithionite-containing sample, suggesting that removal of dithionite did not significantly impair Cu incorporation.

However, XAS and EXAFS spectra reveal structural differences between the NB ± dithionite species. Cu K-edge XAS reveals that the Cu centers remain reduced (Fig. 3a) in the dithionite-free sample, indicating the presence of additional reducing equivalents in the expression condition. However, the 1s→4p transition decreases in intensity and shifts to higher energies, consistent with an increase in the average Cu coordination number^10^. A rising-edge feature at 8980 eV characteristic of four-coordinate Cu(I) is observed in the HERFD-XAS (Fig. S8)^10^. Comparison to the EXAFS of the samples with dithionite reveals similar low-k features, while the higher-k components corresponding to the Cu-Cu interaction are diminished (Fig. 3b). EXAFS fits give Cu-S and Cu-Cu distances nearly identical to samples with dithionite, but with significantly larger σ^2^ for the Cu-Cu component (Table S2 and Fig. S9), suggesting a disordered or potentially misformed site with partial occupancy of the binuclear cluster. The ordered first shell with the Cu-S path suggests that the proportion of Cu that fails to be assembled into the cluster is small, and the limited Cu-N/O contribution to the EXAFS likewise supports minimal adventitious or nonspecific binding of Cu to NB. These data suggest that dithionite may also be responsible for promoting assembly of the Cu site in addition to the [Fe_4_S_4_]^2+^ cluster in NB-protein.

**Figure 3.**
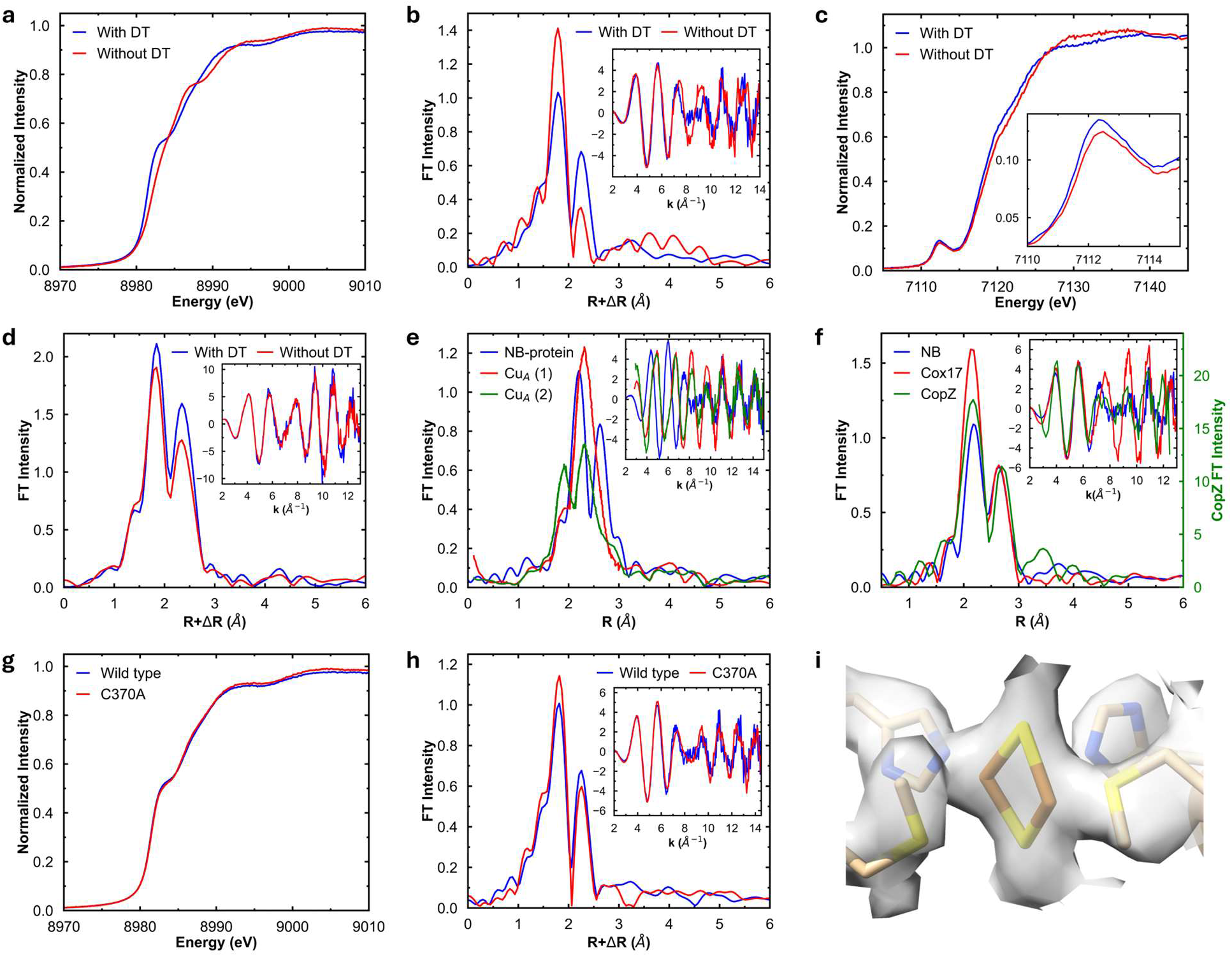
Characterization of sulfur ligands of the Cu-S cluster in NB-protein. **a-b**, Cu K-edge XAS (**a**) and EXAFS/FT (**b**) of NB-protein prepared with and without dithionite. **c-d,** Fe K-edge XAS (**c**) and EXAFS/FT (**d**) of NB-protein prepared with and without dithionite. **e,** Comparison of Cu K-edge EXAFS/FT of NB-protein with fully reduced Cu_A_^33^. EXAFS phase correction was calculated using a Cu scatterer, and E_0_ was set to 8985 eV to be consistent with the literature. **f**, Comparison of Cu K-edge EXAFS/FT of NB-protein with Cu chaperones^35,36^. EXAFS phase correction was calculated using a sulfur scatterer. **g-h,** Cu K-edge XAS (**g**) and EXAFS/FT (**h**) of the C370A mutant of NB-protein. **i,** Carved electron density of the Cu site with the modeled Cu_2_S_2_ cluster, His-404, and Met-408.

Dithionite is commonly used in the preparation of reduced Cu(I) states in metalloenzymes, and no clear literature precedent exists for dithionite or its degradation products coordinating to Cu(I). However, dithionite is known to decompose in solution and release potentially coordinating sulfur-based ligands^27^. Controls were prepared by treating aqueous Cu^2+^ with a partially decomposed dithionite solution (∼80% purity by UV-Vis; Fig. S10a). The Cu K-edge XAS and EXAFS show that dithionite reduces Cu^2+^ to metallic Cu in the absence of the protein environment (Fig. S10b). In the presence of 150 mM NaCl (mimicking the protein buffer composition), a structurally distinct chloride-bound Cu(I) species was formed (Fig. S10c). These results support that dithionite maintains a reducing environment without coordinating Cu. We also note that the measured NB-protein samples had high Cu Kα fluorescence counts, and a comparison of the NB-protein XAS with a Cu foil spectrum shows that the samples are free of metallic or other spurious Cu contamination (Fig. S8).

Omitting dithionite during NB-protein preparation did not lower the Fe-to-protein ratio as assessed by ICP-MS (Table S1). The Fe K-edge XAS and EXAFS of dithionite-containing and dithionite-free NB-protein samples are similar (Figs. 3c and 3d), indicating similar oxidation state and cluster structure, and comparison with reported XAS and EXAFS of [Fe_4_S_4_]^2+^ clusters indicates the presence of a canonical [Fe_4_S_4_]^2+^ cluster ^28^. However, the dithionite-free sample fits with larger σ^2^ values for both Fe-S and Fe-Fe scattering paths (Table S4 and Fig. S11), consistent with a greater disorder. We note that the [Fe_4_S_4_]^2+^ cluster of NB has three cysteine and one aspartate ligands, and its EXAFS beat pattern (k = ∼7 Å^-1^) is slightly different from the four-cysteine-ligated clusters^28^ but similar to the [Fe_4_S_4_] cluster of acetol dehydrogenase (AceDH), which has the same ligand set^29^. The cysteine-to-aspartate substitution has been speculated to destabilize the cluster, making it more susceptible to oxidative damage^30^. Consistent with this, both our data (Fig. 2h and Table S1) and the recent AceDH study show sub-stoichiometric Fe loading, implying partial population of the Fe sites^29^.

### The Cu-S cluster is likely sulfide-bridged and resembles Cu-trafficking sites

The EXAFS-derived Cu-S distance (∼2.26 Å) is unlikely to result from Cu(I)-S(Met-408) coordination and is more consistent with thiolate or sulfide coordination. The structural motif suggested by the EXAFS simulations is reminiscent of the binuclear mixed-valent Cu center in Cu_A_, which has two bridging cysteine ligands and a diamond Cu_2_S_2_ core with a short Cu-N(His) coordination^31^. The EXAFS data for both fully reduced Cu(I)_2_ and mixed-valent Cu(I)Cu(II) forms exhibit short Cu-Cu and Cu-S distances, similar to NB-protein^32,33^, and a comparison of fully reduced Cu_A_ from two organisms (reproduced from ref^33^) and the NB protein is shown in Fig. 3e. The phase and amplitude in the k = ∼2-9 Å^-1^ are distinct and are the average of a short Cu-N(His) and two slightly longer Cu-S distances in Cu_A_, while NB has two strong Cu-S interactions and a weaker 0.5 Cu-N/O coordination. In contrast, the EXAFS data for the NB-protein closely resemble those found in Cu-sensing and Cu-trafficking proteins, many of which form multinuclear Cu(I)-thiolate clusters with trigonal Cu(I) coordination and bridging cysteine ligands. Examples such as COX17 and CopZ show Cu-S distances near 2.26 Å and short Cu-Cu contacts between 2.6 and 2.7 Å (Table S5)^20,34^. Their EXAFS spectra (reproduced from refs^35,36^) display similar beat patterns and two characteristic features in Fourier transforms as NB-protein (Fig. 3f). These similarities suggest that the NB-protein Cu site adopts a local structure more typical of Cu chaperones than Cu_A_, dominated by Cu-S and Cu-Cu interactions. However, this resemblance is limited to the Cu coordination environment. Many cytosolic Cu chaperones are small proteins that bind Cu(I) through Cys-containing motifs^37,38^. Some can bind multiple Cu(I) ions or undergo Cu-dependent oligomerization^39,40^. Thus, NB-protein remains distinct from Cu chaperones in its structure and function.

NB protein contains 28 cysteines dispersed in the sequence without forming CxxC motifs, and none of them are surface-exposed, consistent with the observed lack of nonspecific Cu binding. Cys-370 is the nearest cysteine to the NB-protein Cu site, which is positioned ∼8 Å from the modeled Cu ions in the cryo-EM structure and is part of the β-sheet next to the BchB C-terminal α-helix. Although this distance makes direct coordination unlikely, we observed electron density connecting Cys-370 and Cu in the cryo-EM map (Fig. S12), suggesting that a conformational change in solution might bring Cys-370 closer to the Cu centers. We prepared the C370A mutant of *R. sphaeroides* NB-protein (Fig. S6b). The Cu-to-protein ratio of the C370A mutant is comparable to the wild-type protein (Table S1), suggesting the mutation does not significantly affect Cu incorporation. The Cu K-edge XAS and EXAFS data and the corresponding fits for the C370A mutant (Figs. 3g and 3h) are quantitatively similar to WT (Table S2 and Fig. S13), indicating that Cys-370 does not provide the bridging sulfur ligand.

We re-refined our previously reported WT NB-protein model (PDB 8VQH) against the 2.70 Å-resolution cryo-EM map, incorporating a Cu_2_S_2_ diamond motif with bridging sulfide ligands. The Cu-Cu and Cu-S distances were restrained to the EXAFS-derived values, while the protein ligands were left unrestrained. In the refined model, the cross-shaped electron density at the interface of the C-terminal α-helices well accommodates the Cu_2_S_2_ motif (Fig. 3i). The two Cu-N(His) distances are markedly different (1.96 and 3.56 Å), indicating that only one His-404 engages the cluster, matching the fractional Cu-N coordination number of 0.5 obtained from EXAFS fitting. The Met-408 side chain is less determined, as one Cu-S(Met) distance refines to 1.82 Å, shorter than any chemically reasonable Cu-S bond. Two water molecules can be modeled adjacent to the sulfides but not as Cu ligands (Fig. S14). At ∼2.70 Å resolution, we therefore take the map as evidence for the topology of the site – a Cu_2_S_2_ core with unequal His-404 engagement – rather than for quantitative bond distance assignment, which instead relies on EXAFS.

A [Cu(I)-Fe_4_S_4_] cluster was recently reported in acetol dehydrogenase^29^ (AceDH) with a four-coordinate pendant Cu(I) center that shares one sulfide ligand with the [Fe_4_S_4_] cluster. Its Cu K-edge XAS and EXAFS are distinct from NB-protein (Fig. S15). Particularly, the rising-edge region of the [Cu(I)-Fe_4_S_4_] cluster is characteristic of a four-coordinate Cu(I) site^10^ and significantly different from the three-coordinate NB-protein. Furthermore, a distinct Cu-Fe feature is absent in the Cu K-edge EXAFS data of [Cu(I)-Fe_4_S_4_], unlike the characteristic Cu-Cu feature observed in NB-protein. Together, the cryo-EM, ICP-MS, XAS, EXAFS, and mutagenesis studies presented here argue against Cu binding or incorporation into a [Fe_4_S_4_] cluster in NB-protein.

## Discussion

DPOR is structurally and functionally analogous to nitrogenase, and both display functional asymmetry^7,8,41,42^. Here, we show that the catalytic NB-protein component in DPOR contains an unusual binuclear Cu site at the tetramer interface, conserved in the three purple non-sulfur bacteria examined. Its functional significance is illustrated by the H404A-M408A mutant, which fails to incorporate Cu and lacks catalytic activity, yet retains the heterotetrameric fold and binds L-protein. The loss of activity therefore cannot be explained by gross structural disruption or failure to engage L-protein. However, under turnover conditions, the mutant binds two L-proteins simultaneously (Fig. 1a), in contrast to the asymmetric binding observed for the wild-type (Fig. 1b). This change in binding mode, together with the loss of activity, supports a role for the Cu-binding site in the productive coupling of the two catalytic halves of DPOR.

The bridging ligands of the cluster are sulfur atoms that do not belong to the nearest cysteine or Met-408. Mutation of Cys-370 leaves the EXAFS essentially unchanged, and the 2.26 Å Cu-S distance is too short for Cu(I)-thioether coordination. Chloride is excluded by the compact cluster geometry, as Cu_2_Cl_2_ cores typically have a mean Cu-Cu distance of ∼3.5 Å and wider Cu-Cl-Cu angles than those of the DPOR cluster (Fig. 2c, d). Inorganic sulfide is consistent with the combined structural and spectroscopic evidence. Supporting this assignment, the reducing condition used to stabilize the [Fe_4_S_4_]^2+^ clusters of NB-protein also promotes ordered assembly of the Cu-S cluster (Fig. 3a-d).

Cu-sulfide bonds are rare in biology but not unprecedented. The most thoroughly characterized example is the Cu_Z_ site of nitrous oxide reductase, a tetranuclear µ_4_-sulfide bridged cluster for N_2_O binding and reduction^43^. In other examples, sulfide ligands are shared with other metallocofactors, such as tetrathiomolybdate (the Mo-Cu carbon monoxide dehydrogenase^44^ and the orange protein^45^) and [Fe_4_S_4_] cluster (AceDH^29^), and mononuclear Cu sites are formed. Binuclear Cu(µ-S) clusters are meanwhile well established in synthetic chemistry^46^, but no protein-bound example has been reported. These comparisons make the Cu_2_S_2_ core of the NB-protein, to our knowledge, the first sulfide-bridged binuclear Cu cluster identified in a protein.

Direct confirmation of sulfide ligation faces two obstacles. Incomplete loading of the [Fe_4_S_4_]^2+^ cluster complicates the assignment of acid-labile sulfide to the Cu site (Fig. 2h), and the 28 cysteines and 40 methionines in the tetramer prevent an assignment based on S K-edge XAS. Future X-ray spectroscopy work will focus on replacement of the bridging sulfide with selenide, and selenomethionine incorporation would test the weak Cu-Met interaction proposed from the EXAFS fits. Selenomethionine-substituted NB-protein was previously reported (PDB 3AES), and comparison with the native structure (PDB 3AER) shows a similar side-chain conformation at the position corresponding to Met-408^25^.

The chemistry of the cluster provides further clues to its function. Each Cu is reduced and three-coordinate, and closed-shell 3d^10^ Cu(I) does not provide ligand-field stabilization that would favor a particular geometry. Cu K-edge EXAFS suggests that the DPOR cluster has a similar structure to the Cu(I)-sulfur clusters in Cu-sensing and trafficking proteins, which have Cu-Cu interactions at ∼2.6-2.8 Å (Table S5). A Cu_2_S_2_ core connecting two helices can therefore accommodate small helix displacements by adjusting Cu-Cu distance and Cu-S-Cu angles without breaking bonds. The DPOR cluster also exhibits asymmetric protein ligation: Cu K-edge EXAFS fits yield an average Cu-N/O coordination number of 0.5 per Cu, and the refined cryo-EM model gives unequal Cu-N(His404) distances of 1.96 Å and 3.56 Å. Cu K-edge HERFD-XAS shows two resolved features, corresponding to 1s→4p based transitions, supporting a deviation from ideal trigonal planar Cu coordination. We propose that the cluster geometry is coupled to the conformations of the C-terminal helices, which contact the Pchlide-binding sites and show conformational variability^8,25,47^. In this scenario, Pchlide binding at one active site would be sensed in trans by the helix of the opposing half. The resulting conformational change would be mechanically relayed through the Cu_2_S_2_ cluster to the other helix, which contacts the second active site, thereby coupling the substrate-binding states of the two halves (Fig. 4). Within this model, asymmetric movements of the C-terminal helices could alter the engagement of protein ligands (e.g. His-404) with the Cu_2_S_2_ cluster and thereby contribute to allostery without requiring a change in Cu oxidation state. This, however, also raises the question of why such a potentially reducing dicopper cluster is selected for a conformational switch. Follow-up studies aimed at quantitative oxidation of one or both the Cu sites will address these questions.

**Figure 4.**
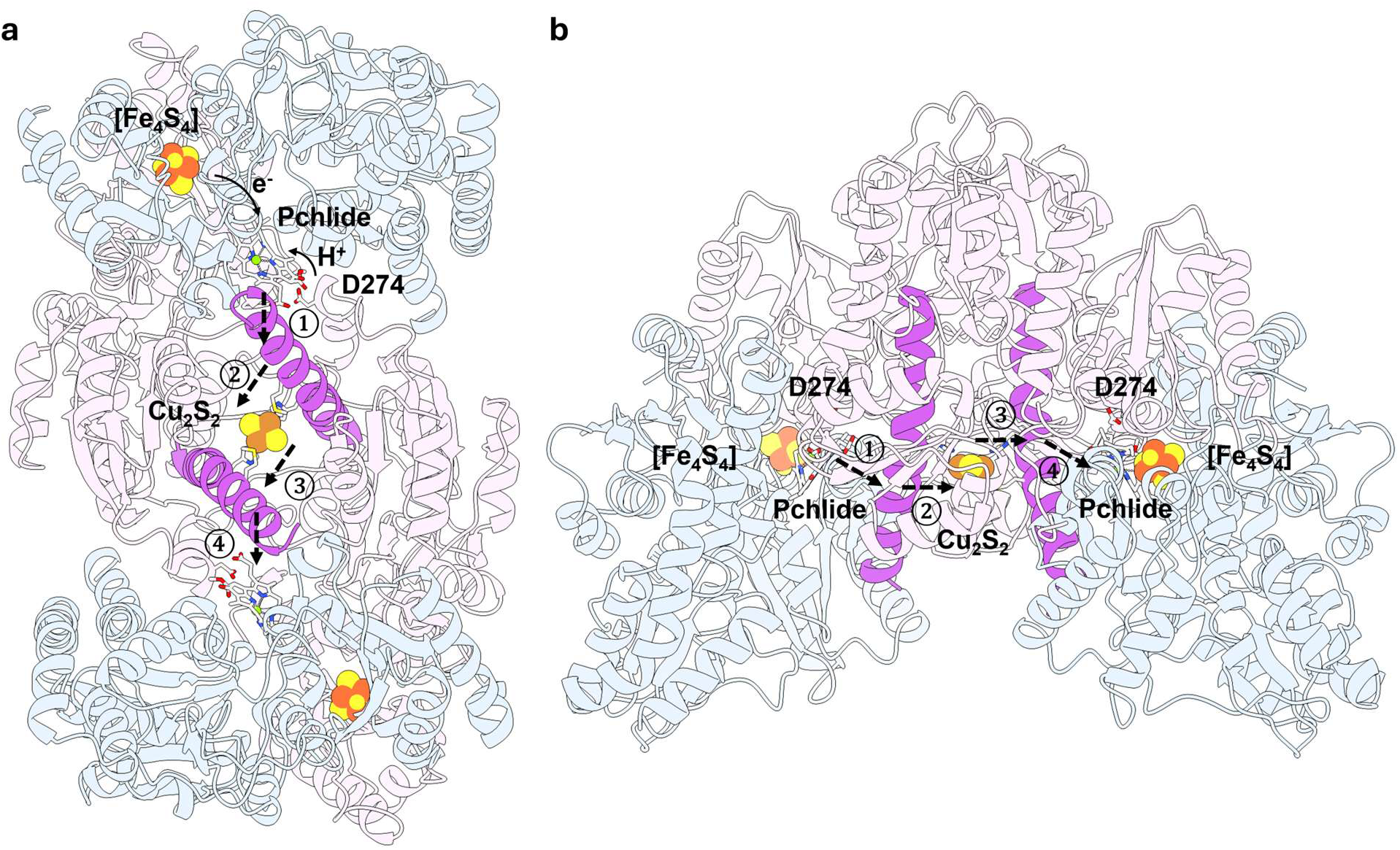
Proposed functional model of NB-protein. Front (**a**) and side view (**b**). The electrons and protons required for Pchlide reduction are provided by the [Fe_4_S_4_] cluster and Asp-274^25^. The binding of Pchlide in one active site is sensed in trans by the contacting C-terminal α-helix of the BchB protein (①). The associated conformational change is relayed mechanically through the Cu-S cluster (②) to the other α-helix (③) and eventually reaches the opposite active site (④).

Nitrogenase achieves analogous cooperativity between its two halves through correlated subunit movements without an interfacial metal site^41^. In DPOR, the [Fe_4_S_4_]^2+^ cluster and Pchlide occupy positions equivalent to the P-cluster and FeMo-cofactor. The Cu_2_S_2_ cluster, however, does not have a counterpart in nitrogenase^6^. Its structural analogs (based on the EXAFS signature) in Cu-trafficking proteins also serve a different biological role. COX17 and CopZ use cysteine-rich motifs at protein surfaces for Cu binding and transfer^48^. By contrast, the DPOR cluster is buried at the center of the heterotetramer and does not use cysteine ligands. It also lies outside the path connecting the [Fe_4_S_4_]^2+^ cluster and Pchlide, making it unlikely to serve as an intermediate in electron transfer. DPOR may instead use a Cu(I)-sulfur core resembling Cu-trafficking sites as an interfacial structural cofactor. The cluster remains intact upon Pchlide binding (Fig. S16), consistent with a persistent structural cofactor that couples the two halves of DPOR during turnover.

The His-404 and Met-408 residues are conserved in the NB-proteins of the three purple non-sulfur bacteria examined and in many other species. However, some homologues lack these residues, including ChlNB from the cyanobacterium *Thermosynechococcus vestitus*. Its crystal structure does not show the Cu cluster reported here at the tetramer interface, and the C-terminal helices are farther apart than those in *R. sphaeroides* (18.4 versus 16.8 Å; Fig. S17)^30^. Whether this protein exhibits functional asymmetry warrants further investigation. Broader comparisons of reactivity across Cu-containing and Cu-free DPORs could clarify how widespread the observed functional asymmetry is and whether it confers a catalytic advantage.

## Methods

### Plasmids

Plasmids for recombinant overproduction of BchL and BchNB were generated as previously described^7,49^. Briefly, the open reading frames for BchB and BchN were PCR amplified from *Rhodobacter sphaeroides* genomic DNA and cloned into the multiple cloning sites 1 and 2 of a pRSF-Duet1 vector, respectively. Similarly, the open reading frames for BchB and BchN were PCR amplified from *Rhodobacter capsulatus and Rhodopseudomonas palustris* genomic DNA but cloned into separate vectors namely pRSF-Duet1 and pCDF vector, respectively. BchB was designed to carry an N-terminal 6X poly-histidine tag followed by a 3C protease cleavage site. Please note that the addition of an affinity tag to either the N- or C-terminus of BchN leads to loss in activity. The BchL open reading frame was similarly PCR amplified and cloned into pRSF-Duet1 with an N-terminal 6X poly-histidine tag followed by a 3C protease cleavage site. The H404A-M408A (double point mutant) and C370A variants were generated using Q5 site-directed mutagenesis (New England Biolabs).

### Protein expression and purification

Wild-type and mutant DPOR component proteins (L-protein and NB-protein) were purified separately using SufFeScient *E. coli* BL21(DE3) cells (PK11466) as previously described with minor modifications^49^. Transformants were grown in 4 to 6 L Luria-Bertani (LB) medium in acid-washed (1 % HCl in water) baffled flasks, supplemented with 1 mM L-cysteine and 1 mM iron (III) citrate. Cultures were grown at 37 °C with shaking at 200 rpm until the optical density of the cultures at 600 nm reached 0.6-0.8. Surfactants should be avoided during cell growth as they lead to degradation/oxidation of the FeS clusters. Protein expression was induced by the addition of Isopropyl β-D-1-thiogalactopyranoside (IPTG) to a final concentration of 0.1 mM. Cultures were grown for an additional 12 h with shaking at 100 rpm at 17 °C. The cells were transferred to centrifuge bottles with rubber gaskets and incubated with 1.7 mM sodium dithionite (unless noted) for 3 h at 17 °C. Care should be taken after this step, and the bottles were briefly opened for the addition of 1 mM CuCl_2_ just an hour before harvesting to avoid exposure to air. Cells were harvested by centrifugation, and the bottles were moved into a Coy Lab anaerobic chamber (glove box) under a 95 % N_2_/5 % H_2_ mixture before decatenation. Further lysis and purification steps were all performed under anoxic conditions and inside the glove box as previously described. All buffers were purged of air under ultrahigh-purity N_2_ on a home-built Schlenk line and stored under N_2_. Cell pellets were resuspended in ∼ 100 ml of Buffer A (100 mM HEPES pH 7.5, 150 mM NaCl, 10 mM MgCl_2_, 10% glycerol, and 1.7 mM of sodium dithionite) supplemented with 3X protease inhibitor cocktail (Sigma Inc.) and 0.1 mg/ml Lysozyme. After incubation at room temperature for 30 min, the cells were lysed by sonication (pulse of 1 sec ON and 3 sec OFF for a total of 3 min inside the glove box), and the lysate was centrifuged at 37,157xg for 60 min at 4 °C. The clarified supernatant was loaded onto a 5 ml Ni^2+^-NTA column (Gold Biotechnology Inc.) equilibrated with Buffer A. Both NB-protein and L-protein complexes were eluted with a step-gradient of imidazole. Fractions containing protein were brown in color and were analyzed on SDS-PAGE for purity (∼ 95% pure). These fractions were pooled and concentrated using an Amicon spin concentrator (30 kDa cutoff) and further fractionated on a S200 HiLoad 26/600 size exclusion chromatography column (Cytiva Healthcare) pre-equilibrated with Buffer A containing no glycerol. Protein-containing fractions were pooled, concentrated, and aliquoted into 1.2 ml cryo-tubes with a gasket-sealed cap (Corning Scientific). Closed tubes with protein were flash frozen in the glove box gas exchanger chamber using liquid nitrogen and stored under liquid nitrogen. Protein concentrations were determined using Bradford reagent with bovine serum albumin (BSA) as the reference.

### Generation of Pchlide

Pchlide was generated from a *Rhodobacter capsulatus* ZY-5 strain harboring a deletion of the BchL gene (a kind gift from Dr. Carl Bauer, Indiana University) and purified as described^50^. Briefly, ZY-5 cells from a frozen glycerol stock were streaked onto an RCV-2/3 agar plate with 0.5 mg/L kanamycin and incubated overnight at 37 °C. The plate was then moved to room temperature, covered with aluminum foil, and allowed to incubate for 2-3 more days. Dark brown colored colonies usually appear in ∼ 2-3 days. A single brown-colored transformant was first grown overnight in 10 ml RCV-2/3 media with 0.5 mg/L kanamycin at 37 °C. This starter culture was then transferred to an aluminum foil-wrapped flask containing 250 mL RCV 2/3 media with 0.5 mg/L kanamycin and grown overnight at 34 °C at 130 RPM. After overnight incubation, the media turned green/dark brown in color, indicating the accumulation of Pchlide. Supernatant containing Pchlide was collected after centrifuging the culture at 4200 RPM for 30 min. At this stage, if more Pchlide is required, the pelleted cells can be resuspended with fresh RCV-2/3 media containing 0.05 mg/L of kanamycin, and the growth can be extended overnight. The supernatant from 5 such collections were pooled, covered in aluminum foil, and stored at 4 °C till further processing. The pooled supernatant solution was spun at 5000 RPM for one hour and filtered through a 0.44 μm Whatman #2 paper filter. The supernatant containing Pchlide was extracted with 1/3 volume of di-ethyl ether in a separating funnel inside a fume hood. The mixture was shaken vigorously and allowed to separate. The aqueous and organic phase layers separate over time. The organic phase contains Pchlide and is subsequently dried using a flow of nitrogen. The dried Pchlide is a dark green/black flaky substance, which is then dissolved in 100 % DMSO. The concentration of Pchlide is determined at 626 nm using ε=30,400 M-1 cm-1. Pchlide was stored at 4 °C in amber colored tubes.

### Preparation of Pchlide-bound double point mutant NB-protein for cryo-EM

Samples were prepared exactly as mentioned previously for the WT NB protein^8^. The poly-histidine affinity tag was first removed from double point mutant NB-protein by treatment with 3 C protease (2 µM protease for 20 µM double point mutant NB-protein) and incubated overnight at RT in the glove box, followed by separation on a S200 Superdex size-exclusion column using buffer A with no glycerol. Fractions containing cleaved double point mutant NB-protein were pooled and concentrated for cryo-EM sample preparation. Double point mutant NB-protein (67 μM tetramer; 134 μM total Pchlide binding sites) was mixed with an excess of Pchlide (520 μM) and passed over a pre-equilibrated SEC column (Superose 6 Increase 5/150 GL; Cytiva Inc.) with Buffer A with no glycerol at room temperature. Peak fractions containing Pchlide-bound double point mutant NB-protein were pooled and concentrated to ∼ 20 μM using an Amicon spin concentrator (30 kDa cutoff). Protein concentration was estimated using a Bradford assay with BSA as reference.

### Cryo-EM data collection

Sample preparation for the turnover complex was performed using a method similar to that previously described for the WT complex^8^. Pchlide-bound Double point mutant NB-protein and L-protein were mixed at a ratio of 1:9 (BchNB:BchL), but in a smaller total volume (10 µl). Glycerol was omitted from the buffer for cryo-EM experiments. The sample was prepared in a sealed glass vial inside the anaerobic chamber. Similarly, a stock of 100 mM ATP in Buffer A was prepared in a separate sealed glass vial. ATP was mixed with the mutant DPOR protein using a Hamilton gas-tight syringe and incubated for 5 min, and immediately applied onto the grid. Cryo-EM samples were prepared on Quantifoil R2/2 (Q350CR2, Electron Microscopy Sciences) holey carbon grids that were plasma cleaned for 60 s using a Solarus 950 (Gatan) with an H2/O2 mixture. Samples were applied onto the grids using a Vitrobot Mark IV (Thermo Fisher Scientific) with the chamber at 4 °C and 100% humidity. To reduce the exposure to the air, a Hamilton gas-tight syringe was used to rapidly apply 3 μl of the sample on the grid from the sealed glass vial. After a 20 s incubation, each grid was blotted for 2 s with a blot force of 0 and immediately plunged into liquid ethane cooled by liquid nitrogen. The total time that each sample spent outside of sealed glass vial during grid preparation and freezing was less than 50 sec. Grids were stored under liquid N2 until data collection. Grids were clipped and loaded into an image-corrected 300 kV Titan Krios G3i cryo-TEM (Thermo Fisher Scientific). Movies were recorded with a Gatan K3 Bioquantum direct electron detector (Gatan Ametek) in an automated fashion using EPU software 2.12.1 (Thermo Fisher Scientific) with a pixel size of 0.865 Å and a nominal defocus range of − 1.0 to − 2.4 µm. Data were acquired with a dose rate of 22 e-/Å2 over a 2.3 s exposure time, yielding a total dose of 53.44 e-/Å2 over 40 fractions. This sample preparation strategy was optimal in achieving consistent uniform thin ice, and no air-water interface issues were encountered. Excellent distribution of diverse and alternate views of the protein in 2D classification were observed for all the samples.

### X-ray absorption spectroscopy

Purified NB-protein samples (in 50 mM HEPES at pH 7.5, 150 mM NaCl, 10 mM MgCl_2_, and 1.7 mM sodium dithionite) were loaded into cuvettes sealed with 25 µm-thick Kapton tape, flash frozen, and stored in liquid nitrogen until measurement. The NB + Pchlide sample was prepared by mixing 150 µM NB with 300 µM Pchlide (this mixture also contains L-protein, which would not bind to NB-protein because no ATP was added). XAS spectra were measured at the Stanford Synchrotron Radiation Lightsource on beamline 7-3. A Si(220) double crystal monochromator, oriented at Φ=0 or 90°, was used for energy selection at the Cu K-edge. The monochromator was detuned by 40% for harmonic rejection. The elastic and inelastic scattering signal from the sample was reduced using a Ni filter and Soller slits. Samples were maintained at 10 K in helium atmosphere during data collection using an Oxford liquid helium cryostat. Spectra were collected in fluorescence mode using a multi-element Canberra Ge detector, and a Cu foil reference spectrum was collected in transmission mode simultaneously for energy calibration with each scan. Multiple spots on each sample were measured (8-12 scans in total) to minimize beam damage, which was not observed during the measurement.

During data processing, channels with large background signals from ice diffraction were removed. The rest of the channels were then averaged in ATHENA^51^. Data were normalized and calibrated by setting the edge energy (first maximum in first derivative) of the Cu foil to 8980.3 eV. The normalized spectra were imported into PySpline^52^ for post-edge background subtraction. EXAFS was extracted by setting E_0_ to 9000 eV and fitting the background using a 4-region spline (with polynomial orders 2, 3, 3, and 3). Fitting of *k*^3^-weighted EXAFS was performed using EXAFSPAK^53^, and the theoretical phase and amplitude of scattering paths were generated using FEFF8 (Figure S3)^54^. The starting geometry of the models was adopted from cryo-EM structures^8^.

High-energy-resolution-fluorescence-detected (HERFD) XANES of NB-protein were measured at beamline 15-2. A Si(311) double crystal monochromator, oriented at Φ=0°, was used for energy selection at the Cu K-edge. Harmonic rejection was provided by a rhodium-coated mirror. Fluorescence signal from the Cu Kα1 line was collected using four Si(444) crystal analyzers and a Vortex silicon drift detector. Samples were maintained at 10 K during data collection using a liquid helium cryostat. Multiple spots on each sample were measured to minimize beam damage, and no beam damage was observed during data collection. A Cu foil reference spectrum was collected in transmission mode simultaneously for energy calibration with each scan. Spectra were exported using PyMca^55^ and then normalized and calibrated in ATHENA^51^.

### Computational methods

DFT and TD-DFT calculations were performed in ORCA 6.1.1^56^. The BP86 functional^57,58^ (with DFT-D4 dispersive correction^59^) and def2-TZVP(-f) basis set^60^ was used for geometry optimization. The auxiliary basis set def2/J was used for Coulomb fitting. Calculations were performed using the unrestricted Kohn-Sham method and tight convergence criteria. Cu K-edge features were simulated with TD-DFT. DFT-optimized geometries were used as input. Relativistic corrections were applied through the Zero-Order Regular Approximation^61^. ZORA-def2-TZVP basis set (ZORA-def2-TZVPP for Cu)^62^ were applied. The auxiliary basis set SARC/J was used for Coulomb fitting. 200 transitions were calculated for each molecule, and each transition was then convoluted into a Gaussian peak with a linewidth of 0.5 eV to be compared with HERFD-XAS. The simulated spectra were shifted by 33.8 eV to align with the experimental spectra.

### Other spectrometry characterizations

UV-Vis spectra of wild-type NB-protein and H404A-M408A mutant of *R. sphaeroides* were collected using a Cary 50 Conc spectrometer at room temperature. The baseline correction was applied using the buffer absorption. The spectra are normalized at the 280 nm and overlaid, as the two proteins have nearly identical sequences and are expected to have similar absorption of the A280 band.

Electron paramagnetic resonance (EPR) spectra of H_2_O_2_-oxidized NB-protein and H404A-M408A mutant of *R. sphaeroides* were collected using a JEOL RE1X EPR spectrometer at 120 K, microwave frequency of 9.188 GHz, modulation amplitude of 0.4 mT, gain of 100, time constant of 0.1 s, and power of 10 mW. The spin quantification was calculated relative to a 1 mM copper acetate standard.

Inductively coupled plasma mass spectrometry (ICP-MS) of NB-protein samples was performed using a Thermo Scientific iCAP RQ ICP-MS instrument. Protein samples were digested using trace metal grade nitric acid.

### Fitting the Cu_2_S_2_ cluster into the cryo-EM map of NB-protein

A ligand restraint file of [Cu_2_S_2_]^2-^ was built using the eLBOW module^63^ of Phenix 2.1^64^. The Cu-S and Cu-Cu distances were restrained to the values derived from EXAFS fitting. The cryo-EM map and model of NB-protein (PDB 8VQH; resolution 2.70 Å) were used for fitting the Cu_2_S_2_ cluster. The two Cu ions, two water molecules, and the alternate location B of Met-408 in the original model were removed. The position of the Cu_2_S_2_ cluster was adjusted using COOT 0.9.8^65^, then the model was further refined using Phenix. ChimeraX^66^ was used for the visualization of the model and density, and the contour level of 0.25 was displayed in Fig. 3i.

## Data availability

Source data are provided as a Source Data file. PDB file deposition number:

## Acknowledgements

This study was supported by funding from the Department of Energy, Office of Basic Energy Sciences DE-SC0020965 (E.A.) and FWP 100593 (R.S.). Use of the Stanford Synchrotron Radiation Lightsource, SLAC National Accelerator Laboratory, is supported by the U.S. Department of Energy, Office of Science, Office of Basic Energy Sciences under Contract No. DE-AC02-76SF00515. The SSRL Structural Molecular Biology Program is supported by the DOE Office of Biological and Environmental Research, and by the National Institutes of Health, National Institute of General Medical Sciences (P30GM133894). The contents of this publication are solely the responsibility of the authors and do not necessarily represent the official views of NIGMS or NIH. R.K. and E.A. also acknowledge generous support from the Doisy Research Fund of the Edward A. Doisy Department of Biochemistry and Molecular Biology at Saint Louis University School of Medicine. We thank Dr. Brock Summers at the Washington University Center for Cellular Imaging for assistance with Cryo-EM data collection.

## Author contributions

Conceptualization: K.Z., R.K., E.A., and R.S. Methodology: K.Z., R.K., E.A., and R.S. Investigation: K.Z., R.K., and M.J.A. Visualization: K.Z. and R.K. Funding acquisition: E.A. and R.S. Supervision: E.A. and R.S. Writing – original draft: K.Z. and R.K. Writing – review & editing: All authors.

**Correspondence** and requests for materials should be addressed to Edwin Antony and Ritimukta Sarangi.

## Reference

1. Calvin, M. The Path of Carbon in Photosynthesis. Science 135, 879–889 (1962).

2. Fujita, Y. Protochlorophyllide reduction: A key step in the greening of plants. Plant and Cell Physiology 37, 411–421 (1996).

3. Fujita, Y. & Bauer, C. E. Reconstitution of light-independent protochlorophyllide reductase from purified Bchl and BchN-BchB subunits: In vitro confirmation of nitrogenase-like features of a bacteriochlorophyll biosynthesis enzyme. Journal of Biological Chemistry 275, 23583–23588 (2000).

4. Masuda, T. & Fujita, Y. Regulation and evolution of chlorophyll metabolism. Photochemical and Photobiological Sciences 7, 1131–1149 (2008).

5. Corless, E. I. et al. The flexible N-terminus of BchL autoinhibits activity through interaction with its [4Fe-4S] cluster and released upon ATP binding. Journal of Biological Chemistry 296, 1–15 (2021).

6. Lago-Maciel, A., Herzog, M. & Rebelein, J. G. Functional insights into the nitrogenase-like enzyme superfamily. Trends in Biochemical Sciences 51, 187–202 (2026).

7. Corless, E. I., Bennett, B. & Antony, E. Substrate recognition induces sequential electron transfer across subunits in the nitrogenase-like DPOR complex. Journal of Biological Chemistry 295, 13630–13639 (2020).

8. Kashyap, R. et al. Cryo-EM captures the coordination of asymmetric electron transfer through a di-copper site in DPOR. Nat Commun 16 3866 (2025).

9. Bedendi, G., Maroni, P. & Milton, R. D. Cooperativity in Enzyme-Substrate Complex Formation in Nitrogenase-like Dark Operative Protochlorophyllide Oxidoreductase (DPOR). Chimia 80, 327–331 (2026).

10. Kau, L. shan, Spira-solomon, D. J., Penner-Hahn, J. E., Hodgson, K. O. & Solomon, E. I. X-ray Absorption Edge Determination of the Oxidation State and Coordination Number of Copper: Application to the Type 3 Site in Rhus vernicifera Laccase and Its Reaction with Oxygen. Journal of the American Chemical Society 109, 6433–6442 (1987).

11. Atkinson, E. R. et al. Tetrakis-[1-methylimidazoline-2(3H)-thione]-μ2-[1-methylimidazoline-2(3H)-thione]-Di-Copper(I) sulphate trihydrate: Preparation, thermal analysis and crystal structure. Inorganica Chimica Acta 100, 285–291 (1985).

12. Seidel, W. W., Arias, M. D. I., Schaffrath, M. & Bergander, K. A neutral tungsten-η2-alkyne-1-thio ligand forming a homoleptic Werner type complex with Cu(I). Dalton Trans. 2053–2054 (2004) doi:10.1039/B404821K.

13. Noda, K. et al. Syntheses and first structural analyses of Cu(I)–PS complexes with bidentate 1,1-diphenyl-1-phospha-4-thiapentane (mtdpp) and quadridentate 5,9-diphenyl-5,9-diphospha-2,12-dithiatridecane (2,3,2-SPPS): successful synthetic route for monomeric [Cu(mtdpp)2]BF4 and dimeric [Cu2(2,3,2-SPPS)2](BF4)2. Inorganica Chimica Acta 357, 526–532 (2004).

14. Liu, Y.-F., Xia, H.-T., Chen, H.-W. & Wang, D.-Q. Bis[(1,10-phenanthroline-κ2N,N′)(2-sulfido-pyridine N-oxide-κ2S,O)copper(II)] tetra-thio-nate trihydrate. Acta Cryst E 62, m3089–m3091 (2006).

15. Zhang, W.-H. et al. Assembly of [(η5-C5Me5)MoS3Cu3]-Supported One-Dimensional Chains with Single, Double, Triple, and Quadruple Strands. Inorg. Chem. 47, 5332–5346 (2008).

16. Brown, K. R. et al. Structures of the cuprous-thiolate clusters of the Mac1 and Ace1 transcriptional activators. Biochemistry 41, 6469–6476 (2002).

17. Eisses, J. F., Stasser, J. P., Ralle, M., Kaplan, J. H. & Blackburn, N. J. Domains I and III of the Human Copper Chaperone for Superoxide Dismutase Interact via a Cysteine-Bridged Dicopper(I) Cluster. Biochemistry 39, 7337–7342 (2000).

18. Chacón, K. N. & Blackburn, N. J. Stable Cu(II) and Cu(I) mononuclear intermediates in the assembly of the CuA center of thermus thermophilus cytochrome oxidase. Journal of the American Chemical Society 134, 16401–16412 (2012).

19. Allen, F. H. The Cambridge Structural Database: a quarter of a million crystal structures and rising. Acta Cryst B 58, 380–388 (2002).

20. Heaton, D. N., George, G. N., Garrison, G. & Winge, D. R. The Mitochondrial Copper Metallochaperone Cox17 Exists as an Oligomeric, Polycopper Complex. Biochemistry 40, 743–751 (2001).

21. Galambas, A. et al. Radical S-adenosylmethionine maquette chemistry: Cx3Cx2C peptide coordinated redox active [4Fe–4S] clusters. J Biol Inorg Chem 24, 793–807 (2019).

22. McKenna, C. E., Gutheil, W. G. & Song, W. A method for preparing analytically pure sodium dithionite. Dithionite quality and observed nitrogenase-specific activities. Biochimica et Biophysica Acta (BBA) - General Subjects 1075, 109–117 (1991).

23. Blackburn, N. J. et al. X-ray absorption edge spectroscopy of copper(I) complexes. Coordination geometry of copper(I) in the reduced forms of copper proteins and their derivatives with carbon monoxide. Inorg. Chem. 28, 1349–1357 (1989).

24. Fritsch, L., Rehsies, P., Barakat, W., Estes, D. P. & Bauer, M. Detection and Characterization of Hydride Ligands in Copper Complexes by Hard X-Ray Spectroscopy. Chemistry - A European Journal 30, 1–8 (2024).

25. Muraki, N. et al. X-ray crystal structure of the light-independent protochlorophyllide reductase. Nature 465, 110–114 (2010).

26. Nomata, J., Ogawa, T., Kitashima, M., Inoue, K. & Fujita, Y. NB-protein (BchN-BchB) of dark-operative protochlorophyllide reductase is the catalytic component containing oxygen-tolerant Fe-S clusters. FEBS Letters 582, 1346–1350 (2008).

27. Telfeyan, K., Migdisov, A. A., Pandey, S., Vesselinov, V. V. & Reimus, P. W. Long-term stability of dithionite in alkaline anaerobic aqueous solution. Applied Geochemistry 101, 160–169 (2019).

28. Mitra, D. et al. Characterization of [4Fe-4S] Cluster Vibrations and Structure in Nitrogenase Fe Protein at Three Oxidation Levels via Combined NRVS, EXAFS, and DFT Analyses. Journal of the American Chemical Society 135, 2530–2543 (2013).

29. Liu, C. et al. An Unprecedented [Cu-4Fe-4S] Cluster in a Methanotroph Acetol Dehydrogenase. 2026.06.11.731722 Preprint at 10.64898/2026.06.11.731722 (2026).

30. Bröcker, M. J. et al. Crystal Structure of the Nitrogenase-like Dark Operative Protochlorophyllide Oxidoreductase Catalytic Complex (ChlN/ChlB)2. Journal of Biological Chemistry 285, 27336–27345 (2010).

31. Solomon, E. I. et al. Copper active sites in biology. Chemical Reviews 114, 3659–3853 (2014).

32. Hay, M. T. et al. Spectroscopic Characterization of an Engineered Purple CuA Center in Azurin. Inorg. Chem. 37, 191–198 (1998).

33. Blackburn, N. J. et al. X-ray absorption studies on the mixed-valence and fully reduced forms of the soluble Cu(A) domains of cytochrome & oxidase. Journal of the American Chemical Society 119, 6135–6143 (1997).

34. Kihlken, M. A., Leech, A. P. & Le Brun, N. E. Copper-mediated dimerization of CopZ, a predicted copper chaperone from Bacillus subtilis. Biochemical Journal 368, 729–739 (2002).

35. Rigby, K., Zhang, L., Cobine, P. A., George, G. N. & Winge, D. R. Characterization of the Cytochrome & Oxidase Assembly Factor Cox19 of *Saccharomyces cerevisiae*. Journal of Biological Chemistry 282, 10233–10242 (2007).

36. Banci, L., Bertini, I., Del Conte, R., Mangani, S. & Meyer-Klaucke, W. X-Ray Absorption and NMR Spectroscopic Studies of CopZ, a Copper Chaperone in Bacillus subtilis: The Coordination Properties of the Copper Ion. Biochemistry 42, 2467–2474 (2003).

37. Huffman, D. L. & O’Halloran, T. V. Function, Structure, and Mechanism of Intracellular Copper Trafficking Proteins. Annual Review of Biochemistry 70, 677–701 (2001).

38. Robinson, N. J. & Winge, D. R. Copper Metallochaperones. Annu. Rev. Biochem. 79, 537–562 (2010).

39. Stasser, J. P., Siluvai, G. S., Barry, A. N. & Blackburn, N. J. A Multinuclear Copper(I) Cluster Forms the Dimerization Interface in Copper-Loaded Human Copper Chaperone for Superoxide Dismutase. Biochemistry 46, 11845–11856 (2007).

40. Hearnshaw, S. et al. A Tetranuclear Cu(I) Cluster in the Metallochaperone Protein CopZ. Biochemistry 48, 9324–9326 (2009).

41. Danyal, K. et al. Negative cooperativity in the nitrogenase Fe protein electron delivery cycle. Proceedings of the National Academy of Sciences of the United States of America 113, E5783–E5791 (2016).

42. Rutledge, H. L., Cook, B. D., Nguyen, H. P. M., Herzik, M. A. & Tezcan, F. A. Structures of the nitrogenase complex prepared under catalytic turnover conditions. Science 377, 865–869 (2022).

43. Rathnayaka, S. C. & Mankad, N. P. Coordination chemistry of the CuZ site in nitrous oxide reductase and its synthetic mimics. Coordination Chemistry Reviews 429, 213718 (2021).

44. Gnida, M., Ferner, R., Gremer, L., Meyer, O. & Meyer-Klaucke, W. A Novel Binuclear [CuSMo] Cluster at the Active Site of Carbon Monoxide Dehydrogenase: Characterization by X-ray Absorption Spectroscopy. Biochemistry 42, 222–230 (2003).

45. George, G. N. et al. A Novel Protein-Bound Copper−Molybdenum Cluster. J. Am. Chem. Soc. 122, 8321–8322 (2000).

46. Delgado, S. et al. Structural diversity of copper complexes with angular and linear dipyridyl ligands. Polyhedron 26, 2817–2828 (2007).

47. Moser, J. et al. Structure of ADP-aluminium fluoride-stabilized protochlorophyllide oxidoreductase complex. Proceedings of the National Academy of Sciences of the United States of America 110, 2094–2098 (2013).

48. Singleton, C., Hearnshaw, S., Zhou, L., Le Brun, N. E. & Hemmings, A. M. Mechanistic insights into Cu(I) cluster transfer between the chaperone CopZ and its cognate Cu(I)-transporting P-type ATPase, CopA. Biochemical Journal 424, 347–356 (2009).

49. Corless, E. I., Mettert, E. L., Kiley, P. J. & Antony, E. Elevated Expression of a Functional Suf Pathway in Escherichia coli BL21(DE3) Enhances Recombinant Production of an Iron-Sulfur Cluster-Containing Protein. J Bacteriol 202, (2020).

50. Corless, E. I. & Antony, E. Methods for Heterologous Overproduction of Fe-S Proteins. in Fe-S Proteins: Methods and Protocols (ed. Dos Santos, P. C.) 69–78 (Springer US, New York, NY, 2021). doi:10.1007/978-1-0716-1605-5_4.

51. Ravel, B. & Newville, M. ATHENA, ARTEMIS, HEPHAESTUS: Data analysis for X-ray absorption spectroscopy using IFEFFIT. Journal of Synchrotron Radiation 12, 537–541 (2005).

52. Tenderholt, A., Hedman, B. & Hodgson, K. O. PySpline: A modern, cross-platform program for the processing of raw averaged XAS edge and EXAFS data. AIP Conference Proceedings 882, 105–107 (2007).

53. George, G. N. & Pickering, I. J. EXAFSPAK: A suite of computer programs for analysis of X-ray absorption spectra. (1995).

54. Ankudinov, A. L., Ravel, B., Rehr, J. J. & Conradson, S. D. Real-space multiple-scattering calculation and interpretation of x-ray-absorption near-edge structure. Phys. Rev. B 58, 7565–7576 (1998).

55. Solé, V. A., Papillon, E., Cotte, M., Walter, P. & Susini, J. A multiplatform code for the analysis of energy-dispersive X-ray fluorescence spectra. Spectrochimica Acta - Part B Atomic Spectroscopy 62, 63–68 (2007).

56. Neese, F., Wennmohs, F., Becker, U. & Riplinger, C. The ORCA quantum chemistry program package. The Journal of chemical physics 152, 224108 (2020).

57. Becke, A. D. Density-functional exchange-energy approximation with correct asymptotic behavior. Physical Review A 38, 3098–3100 (1988).

58. John P. Perdew. Density-functional approximation for the correlation energy of the inhomogeneous electron gas. Physical Review B 33, 8822–24 (1986).

59. Caldeweyher, E. et al. A generally applicable atomic-charge dependent London dispersion correction. The Journal of Chemical Physics 150, (2019).

60. Weigend, F. & Ahlrichs, R. Balanced basis sets of split valence, triple zeta valence and quadruple zeta valence quality for H to Rn: Design and assessment of accuracy. Physical Chemistry Chemical Physics 7 3297 (2005).

61. Van Lenthe, E., Snijders, J. G. & Baerends, E. J. The zero-order regular approximation for relativistic effects: The effect of spin-orbit coupling in closed shell molecules. Journal of Chemical Physics 105, 6505–6516 (1996).

62. Pantazis, D. A. & Neese, F. All-electron basis sets for heavy elements. Wiley Interdisciplinary Reviews: Computational Molecular Science 4, 363–374 (2014).

63. Moriarty, N. W., Grosse-Kunstleve, R. W. & Adams, P. D. electronic Ligand Builder and Optimization Workbench (eLBOW): a tool for ligand coordinate and restraint generation. Acta Cryst D 65, 1074–1080 (2009).

64. Liebschner, D. et al. Macromolecular structure determination using X-rays, neutrons and electrons: recent developments in Phenix. Acta Cryst D 75, 861–877 (2019).

65. Emsley, P., Lohkamp, B., Scott, W. G. & Cowtan, K. Features and development of Coot. Acta Cryst D 66, 486–501 (2010)

66. Pettersen, E. F. et al. UCSF ChimeraX: Structure visualization for researchers, educators, and developers. Protein Science 30, 70–82 (2021).

