## Supplementary Information for "A Binuclear [Cu_2_S_2_] Cluster Supports Functional Asymmetry in Dark-Operative Protochlorophyllide Reductase"

**This PDF file includes:**

Figs. S1 to S17

Tables S1 to S5

References

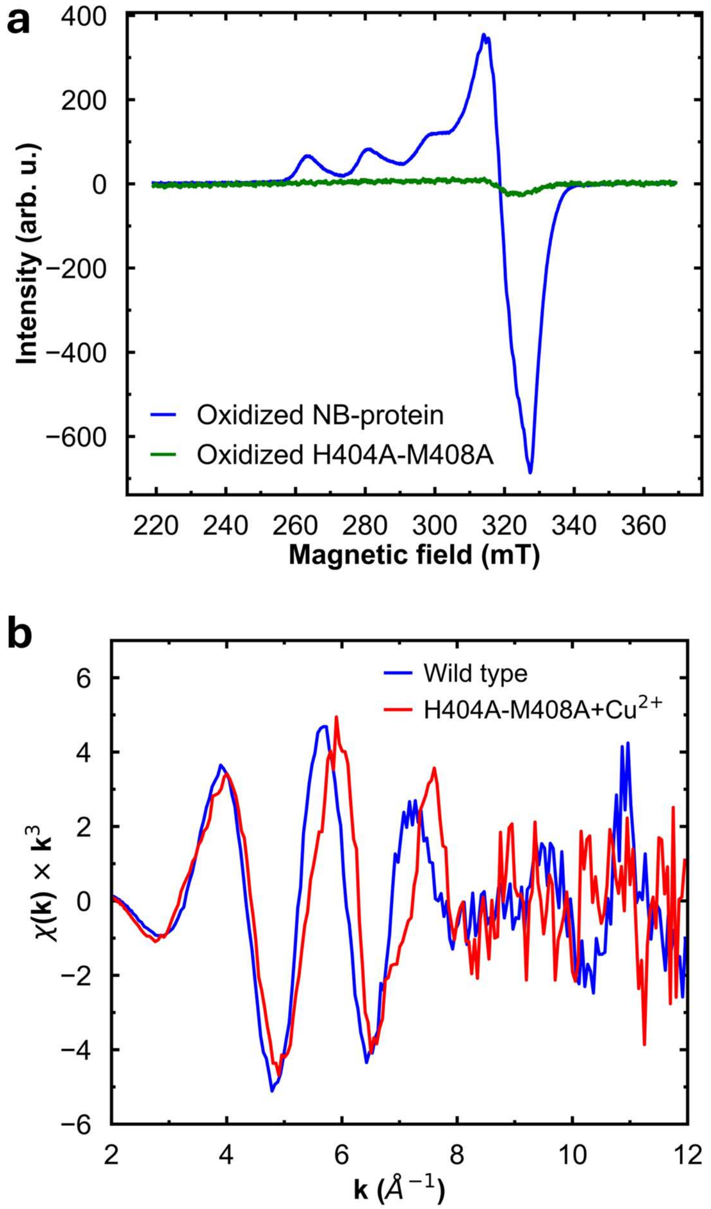


**Figure S1.** Characterization of H404A-M408A mutant. **a,** EPR of $H_2O_2$-oxidized NB-protein and double mutant. **b,** Cu K-edge EXAFS of Cu-supplemented double mutant.

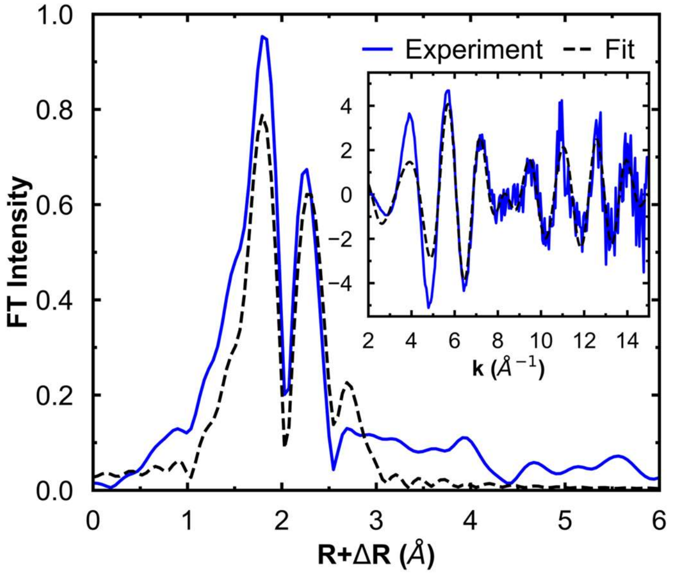


**Figure S2.** Cu K-edge EXAFS and FT of *R. sphaeroides* NB-protein, with fit following the cryo-EM model.

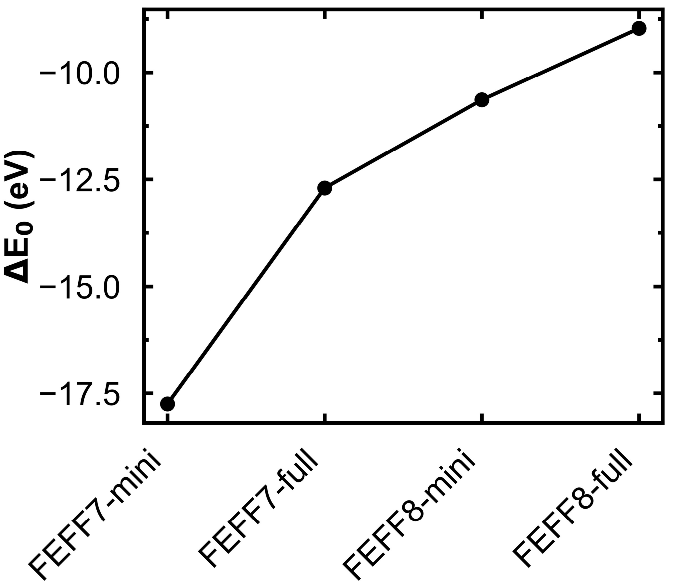


**Figure S3.** $\Delta E_0$ values of Cu K-edge EXAFS fits of *R. sphaeroides* NB-protein, using different versions of FEFF[1,2] and different model sizes for generating the scattering paths. The minimum model (mini) only contains the central Cu and the coordinating atoms, whereas the full model (full) includes all the atoms on H404 and M408 residues. $E_0$ was set to 9000 eV for the fits. The value of $\Delta E_0$ shows a strong dependence on the FEFF version and model size, which could affect the evaluation of the derived structural model. FEFF8 and the full model were selected for fitting all the samples.

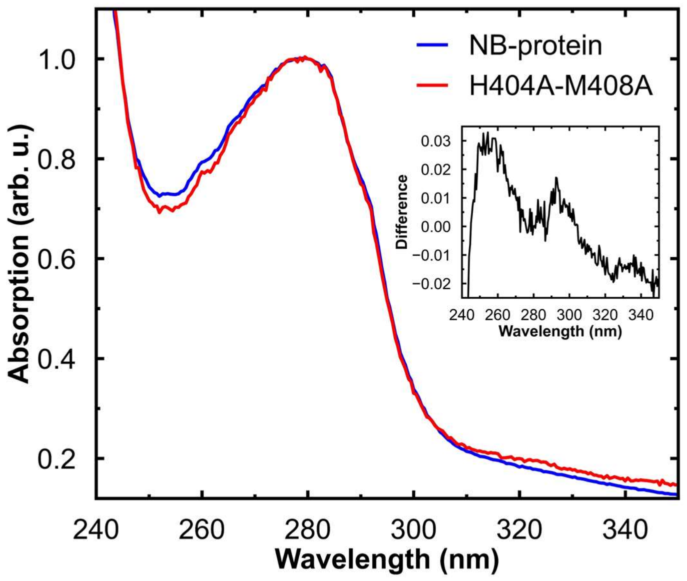


**Figure S4.** UV-Vis absorption spectra of wild-type NB-protein and H404A-M408A double mutant. The inset shows the difference spectrum.

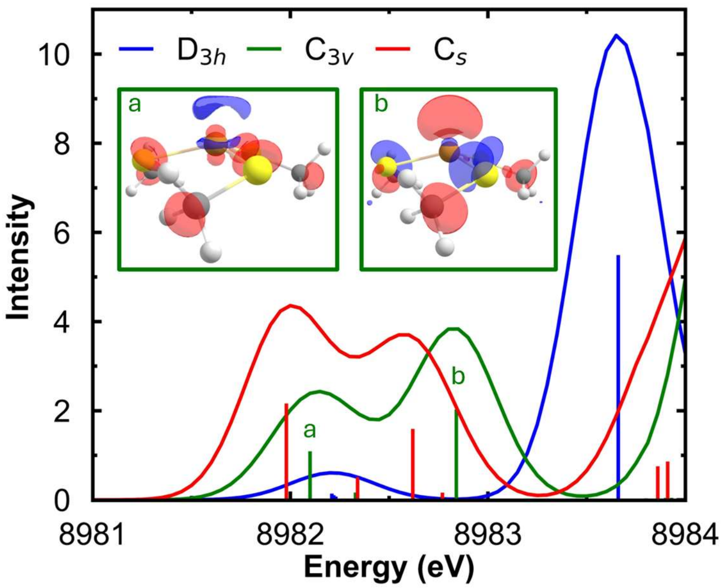


**Figure S5.** TD-DFT simulation of $[Cu(SMe_3)]^{2-}$ model with different degrees of distortion. The inset shows the spin density of two molecular orbitals corresponding to the 1s-4p transition.

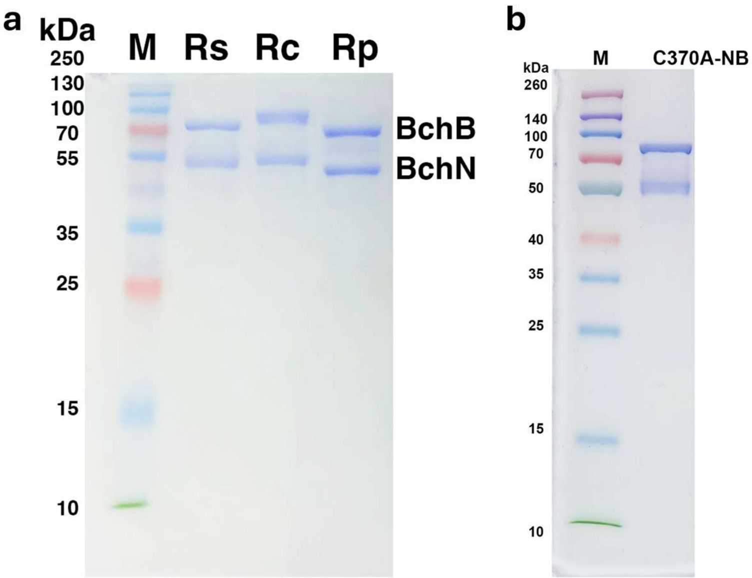


**Figure S6.** SDS-PAGE of purified NB-proteins. (**a**) NB-proteins from *R. sphaeroides* (Rs), *R. capsulatus* (Rc), and *R. palustris* (Rp). (**b**), *R. sphaeroides* C370A NB-protein. Bands for the BchB and BchN subunits are labeled in **a**. M, molecular weight marker (kDa).

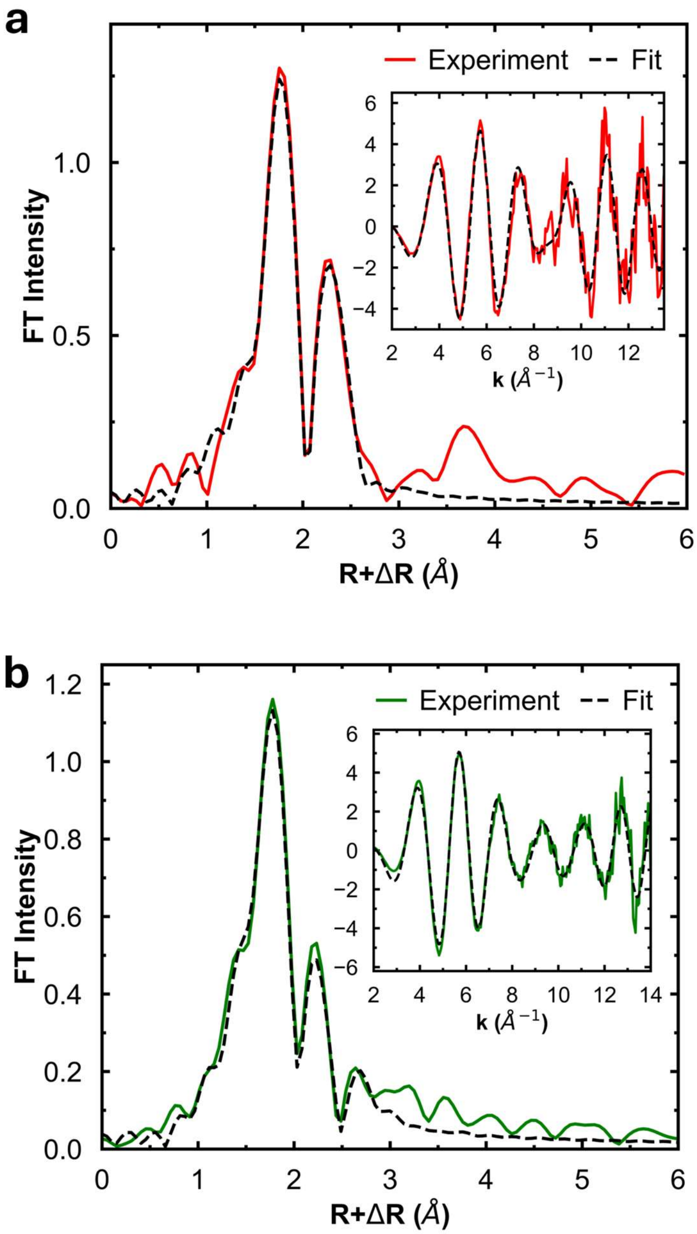


**Figure S7.** Fitting of Cu K-edge EXAFS (inset) and FT of NB-protein of (**a**) *R. capsulatus* and (**b**) *R. palustris*.

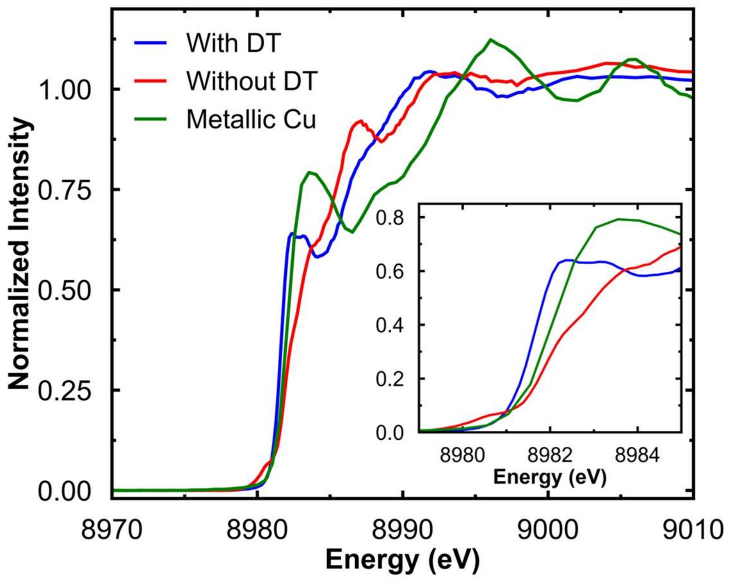


**Figure S8.** Cu K-edge HERFD-XAS of *R. sphaeroides* NB-protein prepared with and without dithionite, overlaid with a Cu foil spectrum[3] for comparison. The inset zooms in on the pre-edge and rising-edge features.

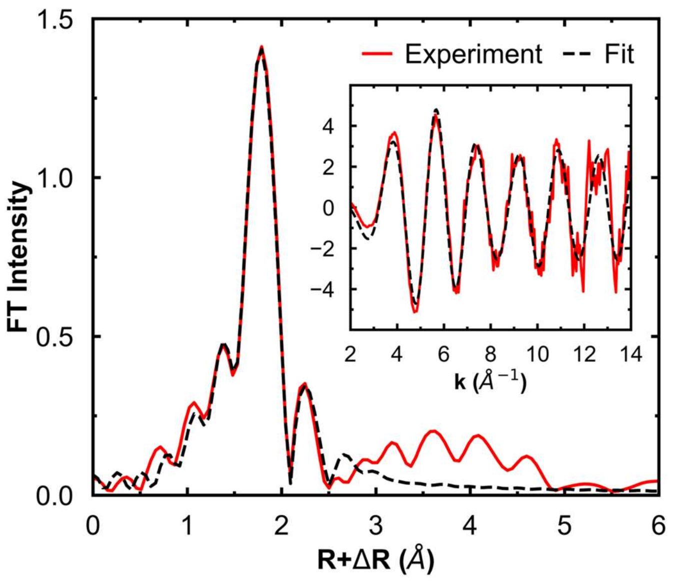


**Figure S9.** FEFF fits of Cu K-edge EXAFS (inset) and FT of *R. sphaeroides* NB-protein, prepared without dithionite.

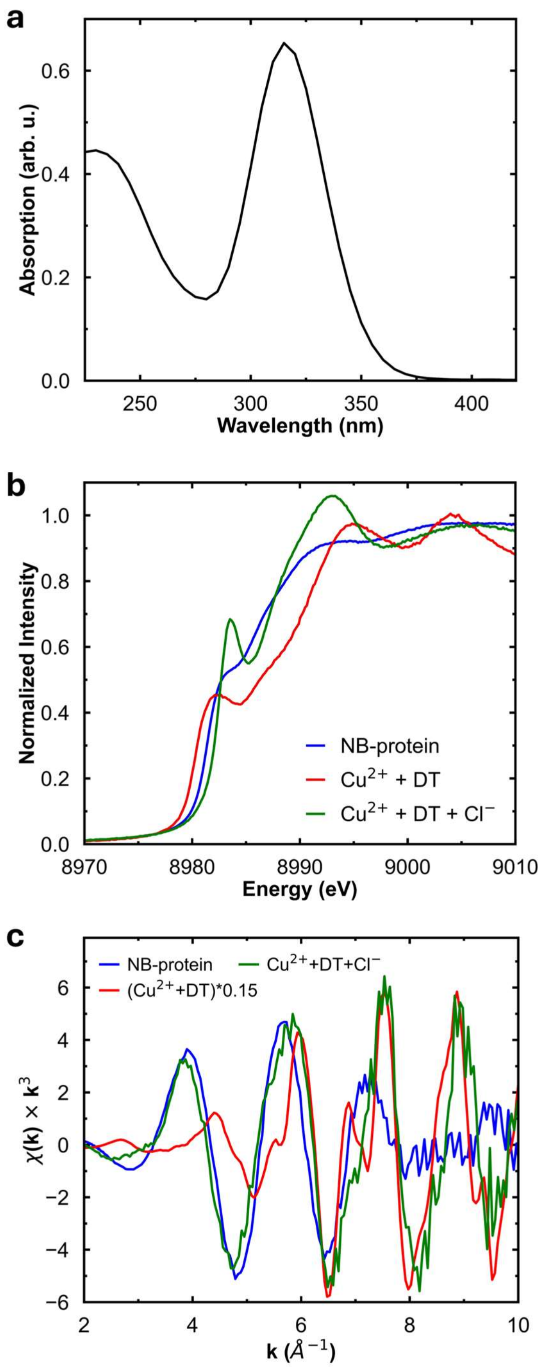


**Figure S10.** Characterization of reaction product between $Cu^{2+}$ and dithionite. **a,** UV-Vis absorption of the dithionite sample. The molar extinction coefficient at 315 nm is 8043 $M^{-1}cm^{-1}$[4], therefore the purity of tested dithionite sample is ~80%. **b-c,** Cu K-edge XAS (**b**) and EXAFS (**c**) of reaction product between $Cu^{2+}$ and dithionite in 50 mM HEPES buffer at pH 7.5, with and without 150 mM NaCl. The red EXAFS spectrum is multiplied by 0.15.

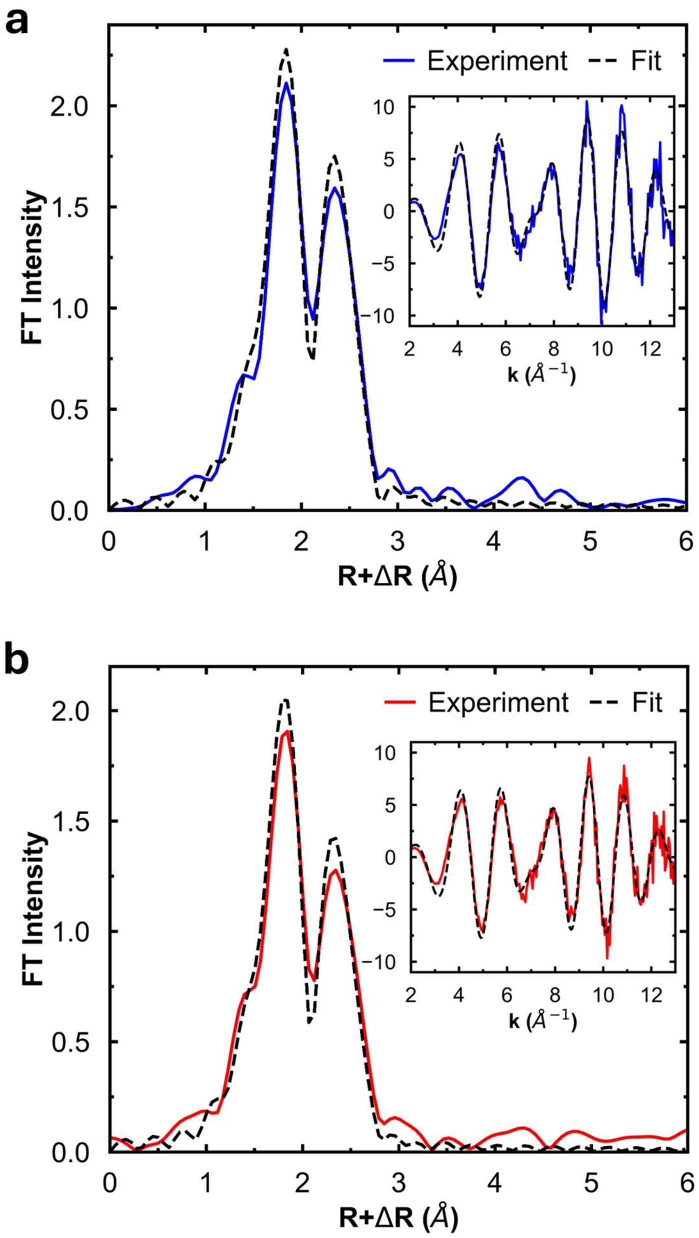


**Figure S11.** FEFF fit of Fe K-edge EXAFS (inset) and FT of *R. sphaeroides* NB-protein, prepared with (**a**) and without (**b**) dithionite.

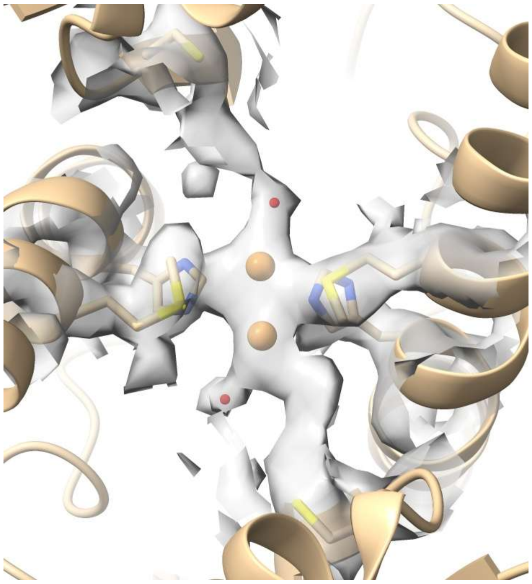

**Figure S12.** Carved electron density of the Cu site with the model of PDB 8VQH[5], showing electron density connecting Cys-370 and Cu ions.

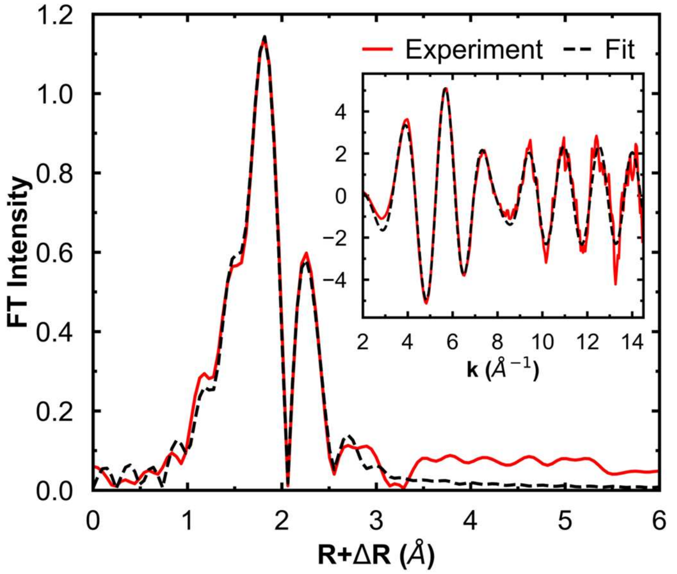


**Figure S13.** Fitting of Cu K-edge EXAFS (inset) and FT of *R. sphaeroides* NB-protein C370A mutant.

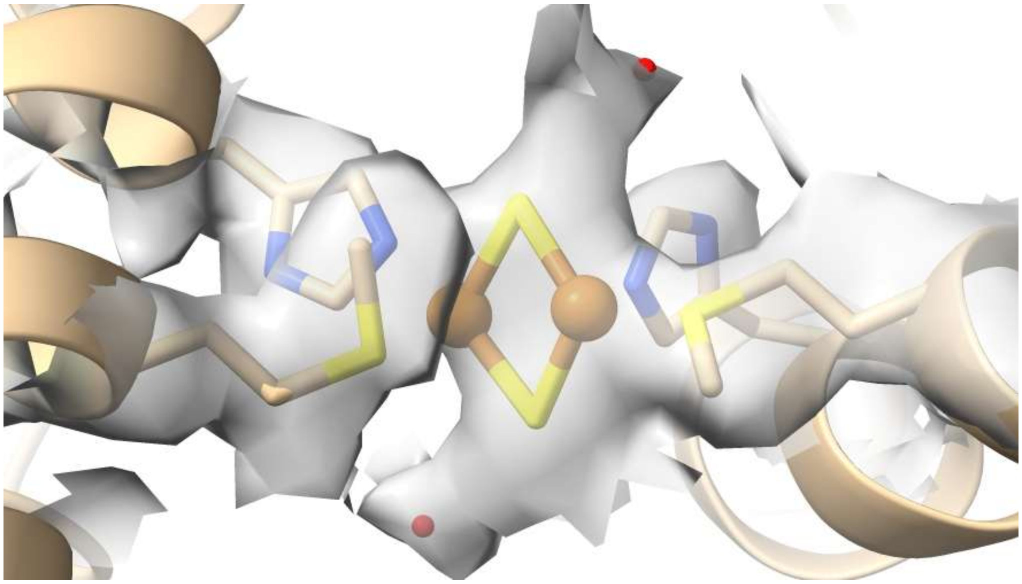

**Figure S14.** Carved electron density of the Cu site (PDB 8VQH[5]) with the modeled $Cu_2S_2$ cluster and two water molecules.

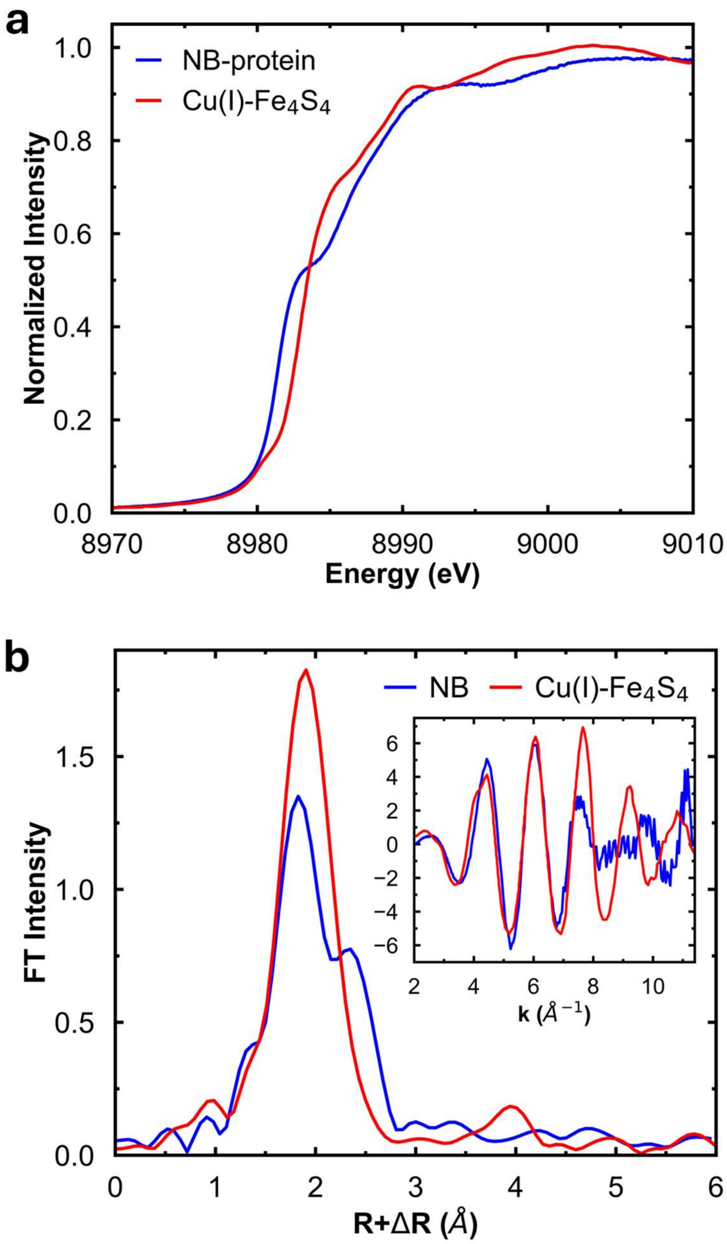


**Figure S15.** Comparison of the NB cluster with the Cu(I)-$Fe_4S_4$ cluster in AceDH[6]. **a,** Cu K-edge XAS. **b,** Cu K-edge EXAFS (inset) and FT.

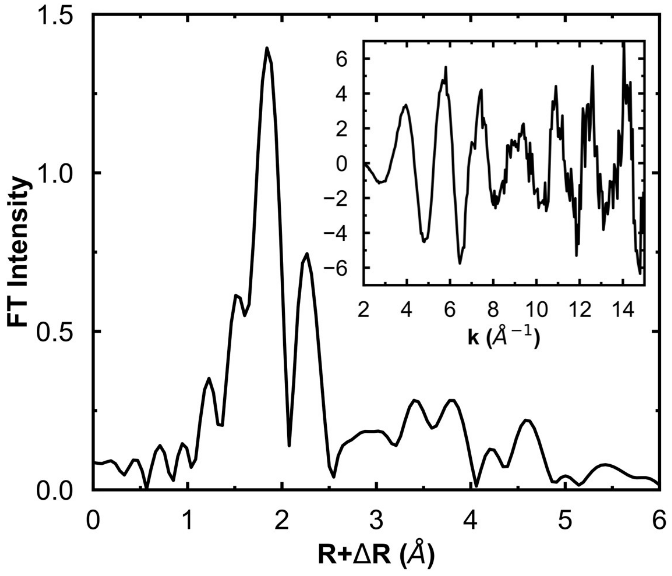


**Figure S16.** Cu K-edge EXAFS and FT of *R. sphaeroides* NB-protein treated with Pchlide.

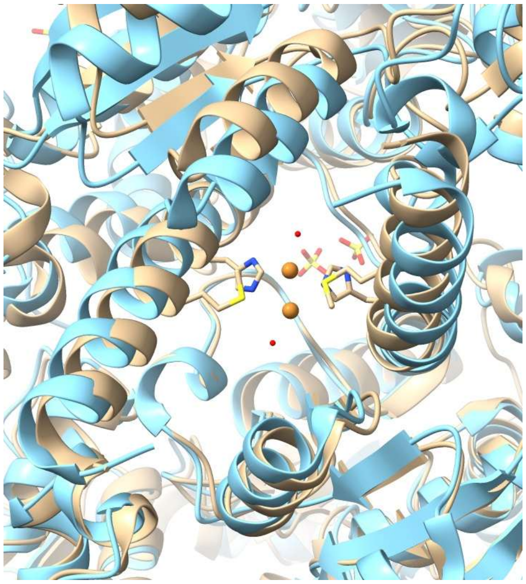

**Figure S17.** Overlay of cryo-EM structure of *R. sphaeroides* BchNB protein (PDB 8VQH[5]; tan) and crystal structure of *Thermosynechococcus vestitus* ChlNB protein (PDB 2XDQ[7]; cyan), which lacks the Cu-binding histidine and methionine residues. The distance between the crossing α-helices is 16.8 Å for BchNB and 18.4 Å for ChlNB. This distance is calculated by fitting one axis to each α-helix (residue 390-419 for *R. sphaeroides* and residue 410-439 for *T. vestitus*) then calculating the distance between the two axes.

**Table S1.** Metal loading of *R. sphaeroides* NB-protein samples determined by ICP-MS. Each column is a separate preparation made by the same protocol.

| | w/ dithionite | w/o dithionite | C370A, w/ dithionite |
|---|---|---|---|
| Cu/NB complex | 1.81 | 2.22 | 2.37 |
| Fe/NB complex | 2.49 | 5.26 | 5.98 |

**Table S2.** FEFF fit results of Cu K-edge EXAFS of NB-protein.

| Molecule | Coordination/Path | *R* (Å) | $\sigma^2$ (Å$^2$)×10$^5$ | $\Delta E_0$ (eV) | *F** |
|---|---|---|---|---|---|
| NB-protein (*R. sphaeroides*; cryo-EM model) | 2 Cu-N/O | 2.16 | 173 | 4.75 | 0.476 |
| | 1 Cu-Cu | 2.68 | 488 | | |
| | 1 Cu-S | 2.97 | 261 | | |
| NB-protein (*R. sphaeroides*; w/o Cu-N path) | 2 Cu-S | 2.26 | 580 | -8.77 | 0.374 |
| | 1 Cu-Cu | 2.62 | 415 | | |
| | 1 Cu-S | 2.89 | 400 | | |
| NB-protein (*R. sphaeroides*; w/ 0.5 Cu-N path) | 0.5 Cu-N | 1.95 | 983 | -8.97 | 0.366 |
| | 2 Cu-S | 2.26 | 571 | | |
| | 1 Cu-Cu | 2.62 | 414 | | |
| | 1 Cu-S | 2.89 | 398 | | |
| NB-protein (*R. sphaeroides*; w/ 1 Cu-N path) | 1 Cu-N | 2.02 | 2200 | -7.90 | 0.365 |
| | 2 Cu-S | 2.27 | 606 | | |
| | 1 Cu-Cu | 2.63 | 399 | | |
| | 1 Cu-S | 2.90 | 354 | | |
| NB-protein (*R. sphaeroides*; w/ 0.5 2$^{nd}$ Cu-S path) | 0.5 Cu-N | 1.96 | 1026 | -8.96 | 0.374 |
| | 2 Cu-S | 2.27 | 581 | | |
| | 1 Cu-Cu | 2.63 | 401 | | |
| | 0.5 Cu-S | 2.90 | 106 | | |
| NB-protein (*R. sphaeroides*; w/o 2$^{nd}$ Cu-S path) | 0.5 Cu-N | 1.92 | 551 | -11.33 | 0.425 |
| | 2 Cu-S | 2.26 | 561 | | |
| | 1 Cu-Cu | 2.63 | 508 | | |
| NB-protein (*R. sphaeroides*; w/ 2 Cu-O path) | 0.5 Cu-N | 1.92 | 634 | -10.26 | 0.378 |
| | 2 Cu-S | 2.26 | 583 | | |
| | 2 Cu-O | 2.67 | 648 | | |
| | 1 Cu-Cu | 2.62 | 43 | | |
| NB-protein (*R. capsulatus*) | 0.5 Cu-N | 1.84 | 764 | -11.91 | 0.381 |
| | 2 Cu-S | 2.24 | 424 | | |
| | 1 Cu-Cu | 2.61 | 494 | | |
| | 1 Cu-S | 2.78 | 1733 | | |
| NB-protein (*R. palustris*) | 0.5 Cu-N | 1.99 | 951 | -9.36 | 0.252 |
| | 2 Cu-S | 2.25 | 498 | | |
| | 1 Cu-Cu | 2.60 | 450 | | |
| | 1 Cu-S | 2.90 | 286 | | |
| H404A-M408A+$Cu^{2+}$ (*R. sphaeroides*) | 2 Cu-N/O | 2.03 | 460 | -1.61 | 0.502 |
| | 1 Cu-S | 2.32 | 252 | | |
| NB-protein (*R. sphaeroides*; w/o dithionite) | 0.5 Cu-N | 1.87 | 276 | -13.06 | 0.304 |
| | 2 Cu-S | 2.25 | 321 | | |
| | 1 Cu-Cu | 2.58 | 643 | | |
| | 1 Cu-S | 2.87 | 432 | | |
| NB-protein (*R. sphaeroides*; C370A mutant) | 0.5 Cu-N | 1.96 | 502 | -8.91 | 0.223 |
| | 2 Cu-S | 2.26 | 429 | | |
| | 1 Cu-Cu | 2.63 | 494 | | |
| | 1 Cu-S | 2.90 | 403 | | |

*The error (*F*) is given by $[\Sigma k^6(\chi_{exptl}-\chi_{calcd})^2/\Sigma k^6\chi_{exptl}^2]^{1/2}$. $S_0^2$ was set to 1.

**Table S3.** MO spin populations of Cu(I) 1s-4p transition features in $[Cu(I)(SCH_3)_3]^{2-}$.

| Symmetry | R(Cu-S) (Å) | MO | Intensity | %4s | %$4p_z$ | %$4p_x$ | %$4p_y$ |
|---|---|---|---|---|---|---|---|
| $D_{3h}$ | 2.303 | 57 | 5.49 | 0.0 | 21.8 | 3.4 | 7.9 |
| $C_{3v}$ | 2.324 | 53 | 1.09 | 39.7 | 6.5 | 0.0 | 0.0 |
| | | 56 | 2.03 | 6.5 | 11.5 | 0.0 | 0.0 |
| $C_s$ | 2.277/2.291/2.880 | 53 | 2.16 | 40.7 | 9.6 | 1.3 | 0.2 |
| | | 54 | 0.51 | 2.3 | 0.5 | 2.0 | 0.2 |
| | | 55 | 1.59 | 3.0 | 9.3 | 0.0 | 0.0 |

**Table S4.** Feff fit results of Fe K-edge EXAFS of NB-protein.

| Molecule | Coordination/Path | $R$ (Å) | $\sigma^2$ (Å$^2$)×10$^5$ | $\Delta E_0$ (eV) | $F$ |
|---|---|---|---|---|---|
| NB-protein (with dithionite) | 4 Fe-S | 2.30 | 437 | -6.86 | 0.263 |
| | 3 Fe-Fe | 2.76 | 529 | | |
| NB-protein (without dithionite) | 4 Fe-S | 2.29 | 514 | -7.57 | 0.288 |
| | 3 Fe-Fe | 2.76 | 662 | | |

**Table S5.** Survey of Cu-S cluster structures in Cu-sensing and trafficking proteins, characterized by Cu K-edge EXAFS.

| Protein | Cluster | Cu coord | R(Cu-S) (Å) | R(Cu-Cu) (Å) | Cu-binding motif | Function | Ref |
|---|---|---|---|---|---|---|---|
| COX11 | $Cu_2S_6$ | 3 | 2.25 | 2.71 | C111/C208/C210 | Cu chaperone | [8,9] |
| COX17 | $Cu_2S_X$ | 3 | 2.26 | 2.70 | CCxC | Cu chaperone | [10–12] |
| COX19 | $Cu_2S_X$ | 3 | 2.26 | 2.70 | Cx(9)C | Cu chaperone | [13] |
| hCCs | $Cu_XS_Y$ | 3 | 2.26 | 2.72 | MTCXXC | Cu chaperone | [14–16] |
| CopZ | $Cu_2S_X$ | 3 | 2.22 | 2.63 | MxCxxC | Cu chaperone | [17,18] |
| CopY | $Cu_2S_4$ | 3 | 2.24 | 2.69 | CxCxxxxCxC | Transcriptional factor | [18,19] |
| Ace1 | $Cu_4S_X$ | 3 | 2.26 | 2.68 | CxxC | Transcriptional factor | [20] |
| Mac1 | $Cu_4S_6$ | 3 | 2.25 | 2.71/2.89 | CxxC | Transcriptional factor | [21] |
| AMT1 | $Cu_4S_X$ | 3 | 2.26 | 2.75 | C-rich domain | Transcriptional factor | [22] |
| Ctr1 | $Cu_4S_6$ | 3 | 2.25 | 2.71 | CxC | Membrane Cu transporter | [23] |
| ATP7B | $Cu_6S_X$ | 3 | 2.22 | 2.73 | MxCxxC | Membrane Cu transporter | [24] |
| dMTF-1 | $Cu_4S_6$ | 3 | 2.26 | 2.70/2.82 | 6 C | Cu sensing | [25] |
| ArsR | $Cu_2S_4$ | 3 | 2.26 | 2.70 | 4 C | Cu sensing | [26] |
| Metallothionein | $Cu_8S_{12}$ | 3 | 2.23 | 2.7/3.9 | 12 C | Cu sequestration | [27,28] |

### Reference


1. Ankudinov, A. & Rehr, J. Relativistic calculations of spin-dependent x-ray-absorption spectra. *Physical Review B - Condensed Matter and Materials Physics* **56**, R1712–R1716 (1997).
2. Ankudinov, A. L., Ravel, B., Rehr, J. J. & Conradson, S. D. Real-space multiple-scattering calculation and interpretation of x-ray-absorption near-edge structure. *Phys. Rev. B* **58**, 7565–7576 (1998).
3. XES data. *MAX IV* https://www.maxiv.lu.se/beamlines-accelerators/beamlines/balder/science-at-balder/experimental-data/xes-data/ (2022).
4. McKenna, C. E., Gutheil, W. G. & Song, W. A method for preparing analytically pure sodium dithionite. Dithionite quality and observed nitrogenase-specific activities. *Biochimica et Biophysica Acta (BBA) - General Subjects* **1075**, 109–117 (1991).
5. Kashyap, R. *et al.* Cryo-EM captures the coordination of asymmetric electron transfer through a di-copper site in DPOR. *Nat Commun* **16**, 3866 (2025).
6. Liu, C. *et al.* An Unprecedented [Cu-4Fe-4S] Cluster in a Methanotroph Acetol Dehydrogenase. 2026.06.11.731722 Preprint at https://doi.org/10.64898/2026.06.11.731722 (2026).
7. Bröcker, M. J. *et al.* Crystal Structure of the Nitrogenase-like Dark Operative Protochlorophyllide Oxidoreductase Catalytic Complex (ChlN/ChlB)2. *Journal of Biological Chemistry* **285**, 27336–27345 (2010).

8. Carr, H. S., George, G. N. & Winge, D. R. Yeast Cox11, a Protein Essential for Cytochrome *c*Oxidase Assembly, Is a Cu(I)-binding Protein. *Journal of Biological Chemistry* **277**, 31237–31242 (2002).
9. Banci, L. *et al.* Solution Structure of Cox11, a Novel Type of β-Immunoglobulin-like Fold Involved in CuB Site Formation of Cytochrome *c* Oxidase*. *Journal of Biological Chemistry* **279**, 34833–34839 (2004).
10. Srinivasan, C., Posewitz, M. C., George, G. N. & Winge, D. R. Characterization of the Copper Chaperone Cox17 of *Saccharomyces cerevisiae*. *Biochemistry* **37**, 7572–7577 (1998).
11. Heaton, D. N., George, G. N., Garrison, G. & Winge, D. R. The Mitochondrial Copper Metallochaperone Cox17 Exists as an Oligomeric, Polycopper Complex. *Biochemistry* **40**, 743–751 (2001).
12. Voronova, A. *et al.* Oxidative switches in functioning of mammalian copper chaperone Cox17. *Biochem J* **408**, 139–148 (2007).
13. Rigby, K., Zhang, L., Cobine, P. A., George, G. N. & Winge, D. R. Characterization of the Cytochrome *c* Oxidase Assembly Factor Cox19 of *Saccharomyces cerevisiae**. *Journal of Biological Chemistry* **282**, 10233–10242 (2007).
14. Eisses, J. F., Stasser, J. P., Ralle, M., Kaplan, J. H. & Blackburn, N. J. Domains I and III of the Human Copper Chaperone for Superoxide Dismutase Interact via a Cysteine-Bridged Dicopper(I) Cluster. *Biochemistry* **39**, 7337–7342 (2000).

15. Stasser, J. P., Siluvai, G. S., Barry, A. N. & Blackburn, N. J. A Multinuclear Copper(I) Cluster Forms the Dimerization Interface in Copper-Loaded Human Copper Chaperone for Superoxide Dismutase. *Biochemistry* **46**, 11845–11856 (2007).

16. Barry, A. N., Clark, K. M., Otoikhian, A., van der Donk, W. A. & Blackburn, N. J. Selenocysteine Positional Variants Reveal Contributions to Copper Binding from Cysteine Residues in Domains 2 and 3 of Human Copper Chaperone for Superoxide Dismutase. *Biochemistry* **47**, 13074–13083 (2008).

17. Banci, L., Bertini, I., Del Conte, R., Mangani, S. & Meyer-Klaucke, W. X-Ray Absorption and NMR Spectroscopic Studies of CopZ, a Copper Chaperone in Bacillus subtilis: The Coordination Properties of the Copper Ion. *Biochemistry* **42**, 2467–2474 (2003).

18. Cobine, P. A. *et al.* Copper Transfer from the Cu(I) Chaperone, CopZ, to the Repressor, Zn(II)CopY: Metal Coordination Environments and Protein Interactions. *Biochemistry* **41**, 5822–5829 (2002).

19. Glauninger, H. *et al.* Metal-dependent allosteric activation and inhibition on the same molecular scaffold: the copper sensor CopY from Streptococcus pneumoniae. *Chemical Science* **9**, 105–118 (2017).

20. Dameron, C. T. *et al.* A copper-thiolate polynuclear cluster in the ACE1 transcription factor. *Proc. Natl. Acad. Sci. U.S.A.* **88**, 6127–6131 (1991).

21. Brown, K. R. *et al.* Structures of the cuprous-thiolate clusters of the Mac1 and Ace1 transcriptional activators. *Biochemistry* **41**, 6469–6476 (2002).

22. Thorvaldsen, J. L. *et al.* Mixed Cu+ and Zn2+ Coordination in the DNA-Binding Domain of the AMT1 Transcription Factor from Candida glabrata. *Biochemistry* **33**, 9566–9577 (1994).

23. Xiao, Z., Loughlin, F., George, G. N., Howlett, G. J. & Wedd, A. G. C-Terminal Domain of the Membrane Copper Transporter Ctr1 from *Saccharomyces cerevisiae* Binds Four Cu(I) Ions as a Cuprous-Thiolate Polynuclear Cluster: Sub-femtomolar Cu(I) Affinity of Three Proteins Involved in Copper Trafficking. *J. Am. Chem. Soc.* **126**, 3081–3090 (2004).

24. Ralle, M., Lutsenko, S. & Blackburn, N. J. Copper transfer to the N-terminal domain of the Wilson disease protein (ATP7B): X-ray absorption spectroscopy of reconstituted and chaperone-loaded metal binding domains and their interaction with exogenous ligands. *Journal of Inorganic Biochemistry* **98**, 765–774 (2004).

25. Chen, X. *et al.* Copper sensing function of Drosophila metal-responsive transcription factor-1 is mediated by a tetranuclear Cu(I) cluster. *Nucleic Acids Res* **36**, 3128–3138 (2008).

26. Liu, T. *et al.* A CuI-Sensing ArsR Family Metal Sensor Protein with a Relaxed Metal Selectivity Profile. *Biochemistry* **47**, 10564–10575 (2008).

27. George, G. N., Byrd, J. & Winge, D. R. X-ray absorption studies of yeast copper metallothionein. *Journal of Biological Chemistry* **263**, 8199–8203 (1988).

28. Smith, T. A., Lerch, K. & Hodgson, K. O. Structural study of the copper sites in metallothionein from Neurospora crassa. *Inorg. Chem.* **25**, 4677–4680 (1986).
